# Inhibiting coronavirus replication in cultured cells by chemical ER stress

**DOI:** 10.1101/2020.08.26.266304

**Authors:** Mohammed Samer Shaban, Christin Müller, Christin Mayr-Buro, Hendrik Weiser, Benadict Vincent Albert, Axel Weber, Uwe Linne, Torsten Hain, Ilya Babayev, Nadja Karl, Nina Hofmann, Stephan Becker, Susanne Herold, M. Lienhard Schmitz, John Ziebuhr, Michael Kracht

**Author notes:** these authors contributed equally to this work (Mohammed Samer Shaban, Christin Müller, Christin Mayr-Buro). **Address correspondence to:** John Ziebuhr, Institute of Medical Virology, Schubertstrasse 81, D-35392 Gießen, Germany, Michael Kracht, Rudolf Buchheim Institute of Pharmacology, Schubertstrasse 81, D-35392 Giessen Germany.

## Abstract

Coronaviruses (CoVs) are important human pathogens for which no specific treatment is available. Here, we provide evidence that pharmacological reprogramming of ER stress pathways can be exploited to suppress CoV replication. We found that the ER stress inducer thapsigargin efficiently inhibits coronavirus (HCoV-229E, MERS-CoV, SARS-CoV-2) replication in different cell types, (partially) restores the virus-induced translational shut-down, and counteracts the CoV-mediated downregulation of IRE1α and the ER chaperone BiP. Proteome-wide data sets revealed specific pathways, protein networks and components that likely mediate the thapsigargin-induced antiviral state, including HERPUD1, an essential factor of ER quality control, and ER-associated protein degradation complexes. The data show that thapsigargin hits a central mechanism required for CoV replication, suggesting that thapsigargin (or derivatives thereof) may be developed into broad-spectrum anti-CoV drugs.

**One Sentence Summary / Running title:** Suppression of coronavirus replication through thapsigargin-regulated ER stress, ERQC / ERAD and metabolic pathways

## Introduction

Coronaviruses are enveloped plus-strand RNA viruses with a broad host range, including humans (de Wit *et al*, 2016; Gorbalenya *et al*, 2020). The four seasonal human CoVs (HCoV-229E, -NL63, - HKU1, -OC43) generally cause a spectrum of (mild) symptoms that are mainly restricted to the upper respiratory tract (Gerna *et al*, 2006; Greenberg, 2011; Jevsnik *et al*, 2012; Nicholson *et al*, 1997). In contrast, the three highly pathogenic CoVs that emerged from animal reservoirs over the past two decades are frequently associated with significant disease burden and mortality in humans. The latter include the severe acute respiratory syndrome (SARS) CoV (Drosten *et al*, 2003; Guan *et al*, 2004; Rota *et al*, 2003), SARS-CoV-2 (Zhou *et al*, 2020; Zhu *et al*, 2020) and Middle East respiratory syndrome CoV (MERS-CoV) (Zaki *et al*, 2012).

The current SARS-CoV-2 pandemic highlights the urgent need to identify new antiviral strategies, including drugs that target the host side (Zhu *et al*., 2020). CoVs impose multiple functional but also structural changes to a wide range of cellular pathways and there is increasing evidence that some of these pathways may be exploited therapeutically (de Wilde *et al*, 2018; Fung & Liu, 2019).

In common with other cellular stress conditions, including infections by diverse pathogens, CoVs are known to activate the NF-κB, JNK and p38 MAPK pathways and to reprogram host cell transcriptomes (Liao *et al*, 2011; Mizutani *et al*, 2005; Poppe *et al*, 2017). In addition, they induce the formation of replicative organelles (ROs), an intracellular network of double-membrane vesicles (DMV) that harbors the viral replication/transcription complexes (RTC) and shields these complexes from recognition by cellular defense mechanisms (Snijder *et al*, 2020). The combination of these and other events leads to cell damage and cell death upon virus budding and release within a few days (de Wilde *et al*., 2018). The virus-induced cellular changes are associated with an activation of the unfolded protein response (UPR) which is evident from a profound transcriptomic endoplasmic reticulum (ER) stress signature, as recently reported for cells infected with HCoV-229E (Poppe *et al*., 2017).

The ER is critically involved in surveying the quality and fidelity of membrane and secreted protein synthesis, as well as the folding, assembly, transport and degradation of these proteins (Wang & Kaufman, 2016). The accumulation of unfolded or misfolded proteins in the ER lumen leads to ER stress and UPR activation, thereby slowing down protein synthesis and increasing the folding capacity of the ER (Karagoz *et al*, 2019). As a result, cellular protein homeostasis can be restored and the cell survives. If this compensatory mechanism fails, ER stress pathways can also switch function and will eventually induce oxidative stress and cell death (Hetz & Papa, 2018; Wang & Kaufman, 2016).

The system relies on three ER membrane-inserted sensors, including the protein kinase R (PKR)-like ER kinase (PERK), inositol-requiring protein 1α (IRE1α) and cyclic AMP-dependent transcription factor 6α (ATF6α). PERK and IRE1α are Ser/Thr kinases whose conserved N termini are oriented towards the ER lumen (Wu *et al*, 2014). In non-stressed cells, the highly abundant major ER chaperone and ER stress sensor binding-immunoglobulin protein BiP (also called 78 kDa glucose-regulated protein, GPR78; heat shock protein family A member 5, HSPA5) binds to PERK and IRE1α, which keeps these two proteins in an inactive monomeric state (Bertolotti *et al*, 2000; Pobre *et al*, 2019). Upon increased binding of BiP to misfolded ER clients, BiP is released from both PERK and IRE1α, resulting in an (indirect) activation of the two kinases by oligomerization and trans(auto)phosphorylation (Carrara *et al*, 2015; Cui *et al*, 2011; Kopp *et al*, 2019).

Active PERK phosphorylates the eukaryotic translation initiation factor 2 (eIF2) subunit α to shut down translation and also activates ATF4, the master transcription factor orchestrating ER stress-induced genes (Han *et al*, 2013; Urra & Hetz, 2017). Phosphorylated IRE1α activates its own RNase domain to generate spliced (s)XBP1 protein, a multifunctional transcriptional regulator responsible for adaptive responses but also cell death (Chen & Brandizzi, 2013). The specific function(s) in this response of yet another ER stress-activated transcription factor, ATF3, is less well understood (Mungrue *et al*, 2009). Generally, the various branches of the UPR act in concert, allowing a multitude of potential outcomes, ranging from the compensation of ER stress and restoration of proteostasis to cell death (Hetz & Papa, 2018).

The activation of ER stress by infectious agents has been widely observed. However, with few exceptions, it remains to be studied how this response is shaped in a microbe-specific manner and whether or not these responses are beneficial or detrimental to the host (Grootjans *et al*, 2016). Moreover, there is a lack of knowledge on CoV-mediated (de)regulation of ER stress components at the protein level. The latter is important because CoVs, in common with many RNA viruses, are known to cause a global shutdown of host protein synthesis (Hilton *et al*, 1986).

Here, we report that CoV infection activates UPR signaling and induces ER stress components at the mRNA level but suppresses them at the protein level. Strikingly, the well-known chemical activator of the UPR, thapsigargin, exerts a profound antiviral effect in the lower nanomolar range on three different CoVs in different cell types. A detailed proteomics analysis reveals multiple thapsigargin-regulated pathways and a network of proteins that are suppressed by CoV but (re)activated by chemically stressed infected cells. These results reveal new insight into central factors required for CoV replication and open new avenues for targeted CoV antivirals.

## Results

To investigate how CoVs modulate ER stress components at the mRNA compared to the protein level, we determined the expression levels of 166 components of the ER stress pathway KEGG 04141 “protein processing in endoplasmic reticulum” in human HuH7 liver cells, a commonly used cellular model for CoV replication, in response to infections with HCoV-229E and MERS-CoV, respectively. For untreated HuH7 cells, we obtained mRNA (by RNA-seq) and protein (by LC-MS/MS) expression data for 119 components which revealed a positive correlation between mRNA and protein abundancies **(Fig. 1A, upper graph)**. However, in cell lysates obtained at 24 h post infection (p.i.), this effect was largely lost **(Fig. 1A, middle and lower graph)**. Pearson correlation matrix confirmed a progressive loss of correlation between mRNA levels and protein levels for this pathway over a time course from 3 h to 24 h p.i. **(Fig. 1B)**. Thus, out of 37 (for HCoV-229E) or 56 (for MERS-CoV) ER stress factors that were found to be regulated at the mRNA level, only a few remained (down)regulated at the protein level at late time points **(Fig. 1C, Fig. S1)**.

**Fig. 1.**
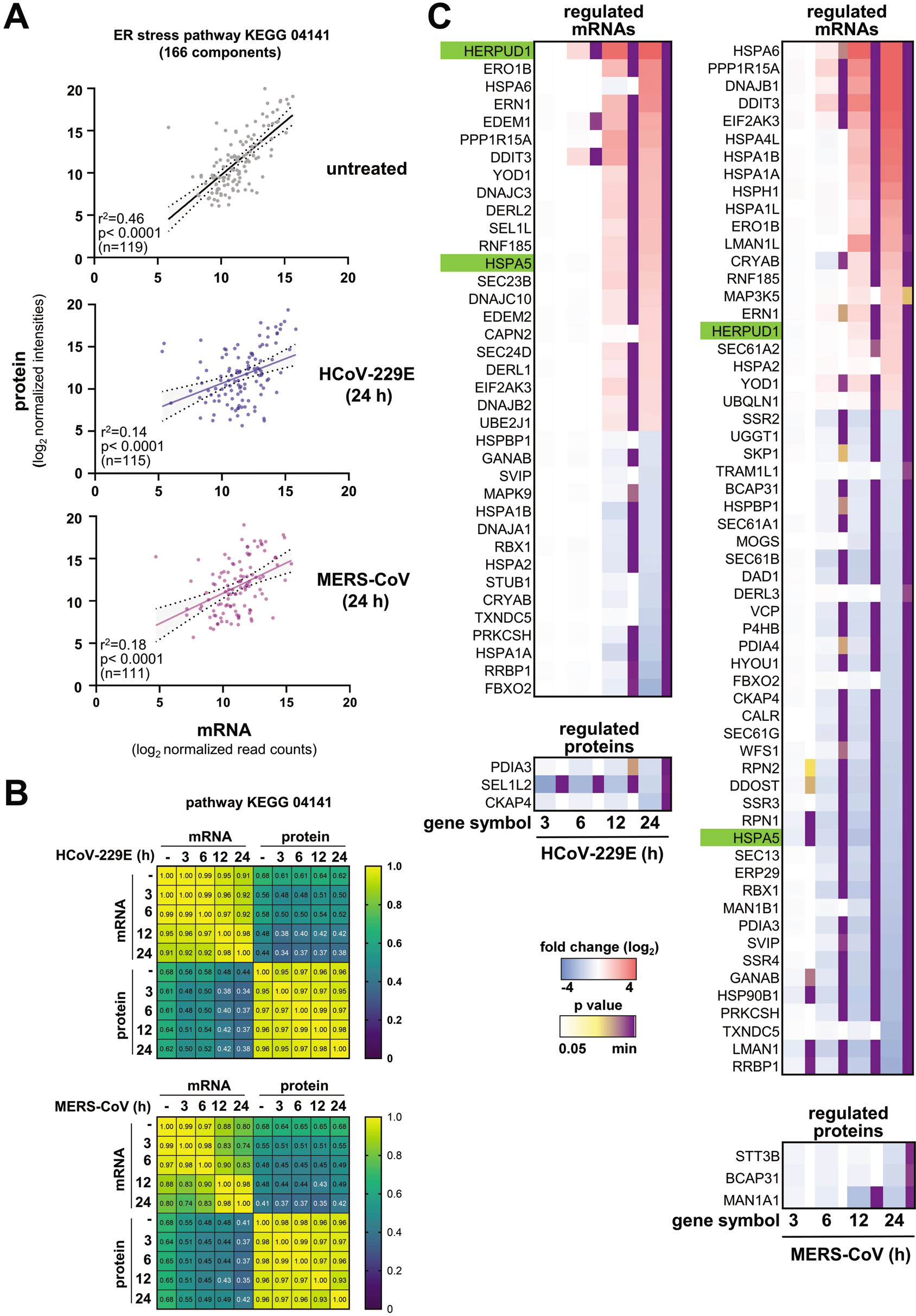
CoVs uncouple mRNA and protein levels of ER stress components in infected cells. HuH7 cells were left untreated or were infected with HCoV-229E or MERS-CoV (MOI=1) for 3, 6, 12, or 24 h. Transcriptomic data (by RNA-seq, n=2) and proteomic data (by LC-MS/MS, n=2, three technical replicates per sample) were derived from samples obtained at the indicated time points p.i. and subsequently used to extract expression values for the KEGG pathway 04141 “protein processing in endoplasmic reticulum”. (A) Scatter plots show mean normalized protein / mRNA expression values for each component, fitted linear regression lines, confidence intervals and correlation coefficients for mock-infected HuH7 cells and HuH7 cells infected for 24 h. (B) Correlation matrix of Pearson’s r across all conditions. All p values are given in supplementary Table S1. (C) The heatmap shows mean ratio values of differentially expressed mRNAs or proteins based on significant differences (fold change ≥ 2, p ≤ 0.01) calculated from the two biological replicates. See also Fig. S1 and Table S1.

To determine the functional consequences of this opposing regulation at the mRNA and protein levels in CoV-infected cells, we focused on HCoV-229E and assessed key regulatory features of the ER stress pathway as shown in **Fig. 2A**. As a reference, we included samples from cells exposed to thapsigargin, a compound that has been widely used to study prototypically activated ER stress mechanistically (Bertolotti *et al*., 2000; Oslowski & Urano, 2011). This setup included experiments, in which thapsigargin and virus were added simultaneously to the cell culture medium (followed by a further incubation for 24 h) or thapsigargin was added to the cells at 8 h p.i. for 16 h **(Fig. 2B)**.

**Fig. 2.**
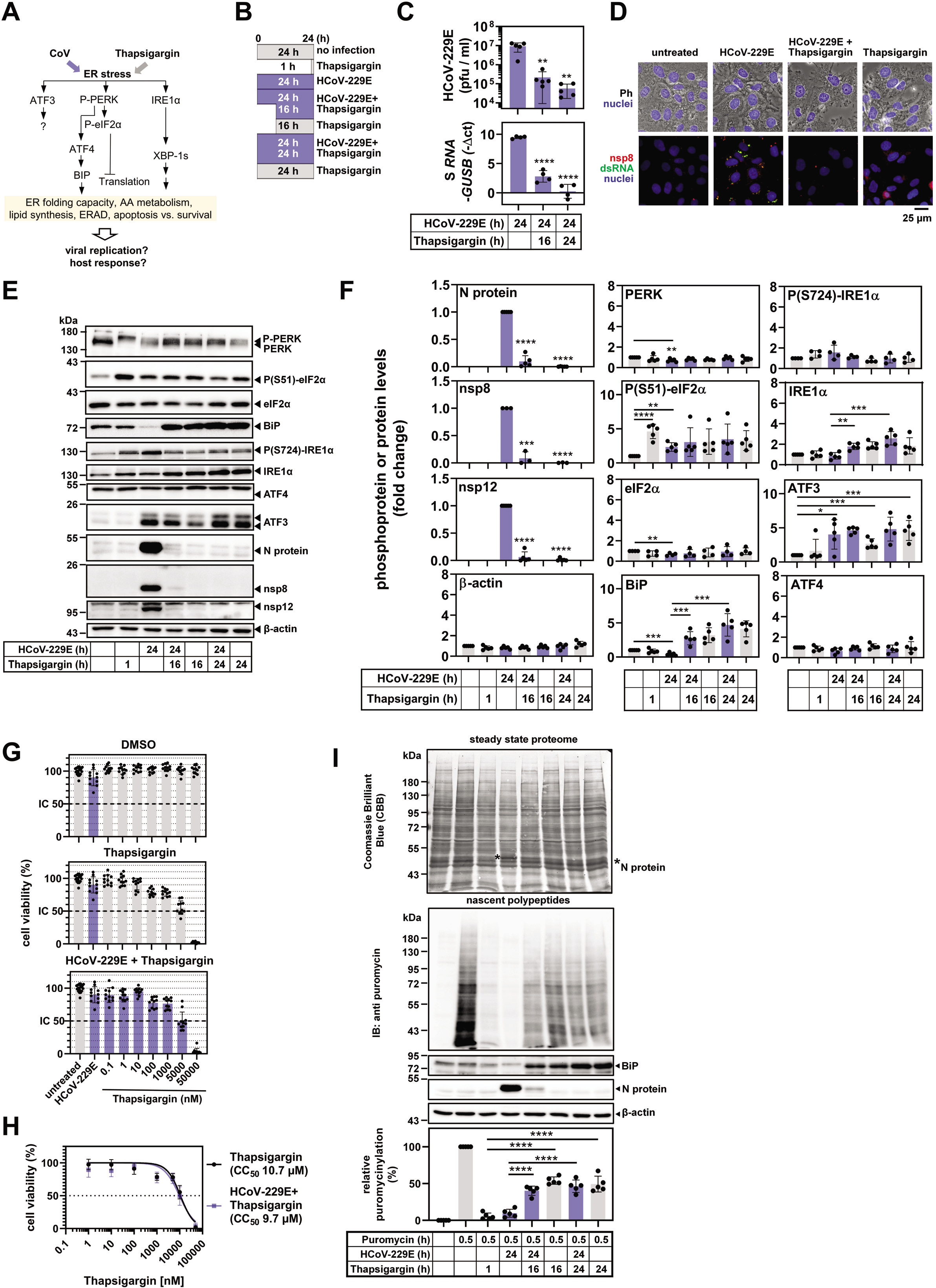
Thapsigargin inhibits HCoV-229E replication, counteracts BiP downregulation and restores protein biosynthesis in virus-infected cells. (A) Schematic overview of parameters used to monitor virus- and thapsigargin-mediated ER stress. (B) Schematic presentation of HCoV-229E infection of cells and/or treatment with thapsigargin for different periods of time as applied in this study. (C) HuH7 cells were left untreated or infected with HCoV-229E (MOI=1) for 24 h and treated with thapsigargin (1 μM) according to (B). Supernatants and RNA-isolated from the cell pellets were used to determine viral titers (upper graphs) and expression of HCoV-229E S gene-containing RNA (lower graphs). (D) Representative fluorescence microscopy images showing subcellular HCoV-229E replication sites (at 24 h p.i.) identified by nsp8- or double-strand RNA-specific antibodies in the presence or absence of thapsigargin. Cells were infected with virus and treated with thapsigargin (1 μM) for 24 h. (E) Total cell extracts from HuH7 cells infected with HCoV-229E (MOI=1) and treated with thapsigargin (1 μM) as shown in (B) were analyzed for the expression/modification of the indicated host cell and viral proteins by immunoblotting. (F) Quantification of immunoblot data from multiple experiments. (G) HuH7 cells were left untreated, or were infected with HCoV-229E (MOI=1) in the presence / absence of increasing concentrations of thapsigargin or DMSO as solvent control. After 24 h, cell viability was assessed by MTS assay. (H) Data from (G) were used to compute half maximal cellular cytotoxicity (CC_50_) concentrations of thapsigargin. (I) HuH7 cells were treated and infected according to (B). 30 min before harvesting, puromycin (3 μM) was added to the cells as indicated. Total cell extracts were analyzed by immunoblotting. The blot membrane was stained with CBB to assess the steady state proteomes and then hybridized with anti-puromycin antibodies to detect *de novo* synthesized nascent polypeptides. The upper graphs show representative images and the lower graph shows the quantification of multiple replicates. Puromycin signals of each lane were normalized to the corresponding CBB staining and were background corrected by subtracting signals of samples in which puromycin had been omitted. Data points show values from independent biological replicates, error bars show s.d.. Asterisks indicate p values (*p ≤ 0.05, **p ≤ 0.01, ***p ≤ 0.001, **** p ≤ 0.0001) obtained by two-tailed unpaired t-tests.

The presence of thapsigargin in the growth medium resulted in a major drop in viral titers by more than 150-fold (from 9.18*10^6^ to 5.7*10^4^ pfu / ml) which was paralleled by reduced amounts of viral RNA isolated from thapsigargin-treated, HCoV-229E-infected infected cells at 24 h p.i. **(Fig. 2C)**. Immunofluorescence analysis of HCoV-229E-infected cells treated with thapsigargin confirmed the impaired formation of functional viral replication/transcription complexes (RTCs) as shown by the reduced levels of both double-stranded RNA (an intermediate of viral RNA replication) and nonstructural protein (nsp) 8 (an essential part of the viral RTC) **(Fig. 2D)**.

A strong suppression of viral replication was also demonstrated by the reduced protein levels observed for the nucleocapsid (N) protein (a major coronavirus structural protein) as well as nonstructural proteins (nsp) 8 and 12, both of which representing essential components of the viral replication complex (Snijder *et al*, 2016) **(Fig. 2E, F)**. In all cases, the antiviral effect of thapsigargin remained readily detectable when the compound was added at 8 h p.i, suggesting that it does not prevent viral entry but rather suppresses intracellular pathways required for efficient RNA replication and/or particle formation and release or activates unknown antiviral effector systems **(Fig. 2C-F)**.

Next, we investigated ER stress signaling under these conditions. Both virus and thapsigargin were confirmed to activate the PERK branch of ER stress **(Fig. 2E, F)**, as shown by the retarded mobility of PERK in SDS gels (indicating multisite phosphorylation) and by phosphorylation of the PERK substrate eIF2α at Ser51 **(Fig. 2E, F)**. Unlike thapsigargin treatment, HCoV-229E infection led to a weak but significant decrease of PERK (mean 71±15 %) and eIF2α (mean 67±13 %) levels compared to the controls. Infection also caused an approximately twofold (mean 42±22 %) reduction in BiP expression **(Fig. 2E, F)**. In contrast, long-term thapsigargin treatment (for 16 h or 24 h) caused a 3–4-fold increase in BiP expression, also in HCoV-229E-infected cells, thus reversing the suppression by viral infection (**Fig. 2E, F)**. Similarly, thapsigargin treatment for 16 h or 24 h caused a 1.5–2-fold increase in IRE1α expression (but not phosphorylation), again also in infected cells (**Fig. 2E, F)**. In this set of experiments, ATF3 proved to be the only protein that was induced by virus alone (**Fig. 2E, F)**, while the expression levels of ATF4 remained largely unchanged **(Fig. 2E, F)**.

These data show that both CoV infection and chemicals like thapsigargin activate ER stress through the same proximal PERK pathway, although they affect downstream cellular outcomes differentially. The restoration of BiP and IRE1α levels by long-term thapsigargin treatment further suggests that the CoV-induced block of inducible host factors is not irreversible and can be reprogrammed by a (presumably protective) thapsigargin-mediated response. Our comparative analyses of viral replication and host response lead us to conclude that chemically and virus-induced forms of ER stress, although proceeding through the same core PERK pathway, do not simply potentiate each other but rather (somewhat counterintuitively) counteract each other.

To explore a potential pharmacological exploitation of this effect, we assessed the cytotoxicity of the combined thapsigargin treatment and virus infection, because both conditions are known to promote cell death.

At 24 h p.i., cell viability of HCoV-229E-infected HuH7 cells was only marginally reduced (mean 90.02±12.32 %) **(Fig. 2G, upper graph)**. After 24 h of incubation, thapsigargin decreased cell viability in a dose-dependent manner with a CC_50_ of 10.7 μM in line with previous reports **(Fig. 2G, middle graph, Fig. 2H)** (Sehgal *et al*, 2017; Tombal *et al*, 2000). The combination of thapsigargin and HCoV-229E infection did not cause additional cytotoxicity as shown by a nearly identical CC_50_ of 9.7 μM **(lower graph, Fig. 2G, Fig. 2H)**. At 1 μM thapsigargin, i.e., a concentration shown to completely abolish viral protein translation and replication (see above), the cell viability of cells infected with HCoV-229E and treated with thapsigargin was 76.6±7.9 %, suggesting that thapsigargin exerts its antiviral effects at concentrations well below its cytotoxic concentrations **(Fig. 2G, H)**.

To further characterize the metabolic state of the cells under the conditions used in these experiments, we investigated protein *de novo* synthesis. Newly produced proteins were quantified by *in vivo* puromycinylation tagging of nascent protein chains followed by immunoblotting using anti-puromycin antibodies. HCoV-229E was found to shut down protein biosynthesis by 90.3±5.4% while 1 h of thapsigargin treatment led to a shutdown by 94.3±4.3% **(Fig. 2I)**. However, in infected cells, the simultaneous or delayed addition of thapsigargin restored (or rescued) protein biosynthesis to approximately 50 % of the level observed in untreated cells **(Fig. 2I)**. These data demonstrate that, although both viral infection and thapsigargin treatment (individually) induce ER stress and cause a translational shut-down, their combination shows no additive harmful effects to the cells. On the contrary, their combination appears to have opposing effects that result in a partial restoration of the cellular metabolic capacity while retaining a profound antiviral effect.

We next assessed if these effects were cell-type or virus-specific. In line with the results described above, the antiviral effects of thapsigargin, the reconstitution of the BiP and IRE1α levels and the lack of additional cytotoxicity in infected cells could be confirmed for diploid MRC-5 embryonic lung fibroblasts infected with HCoV-229E **(Fig. 3A-D)**, as well as for HuH7 cells infected with MERS-CoV and Vero E6 African green monkey kidney epithelial cells infected with SARS-CoV-2 **(Fig. 3E-K, Fig. S2A, B)**. MERS-CoV and SARS-CoV-2 replication were suppressed by thapsigargin with an effective concentration (EC_50_) (as judged by virus titration) of 4.8 nM and 260 nM, respectively **(Fig. 3I, J)**, while the CC_50_ for thapsigargin in Vero E6 cells was 17.22 μM **(Fig. 3K)**, resulting in selectivity indices (SI, CC_50_/EC_50_) of 66.2 for SARS-CoV-2 and 2229.2 for MERS-CoV, respectively.

**Fig. 3.**
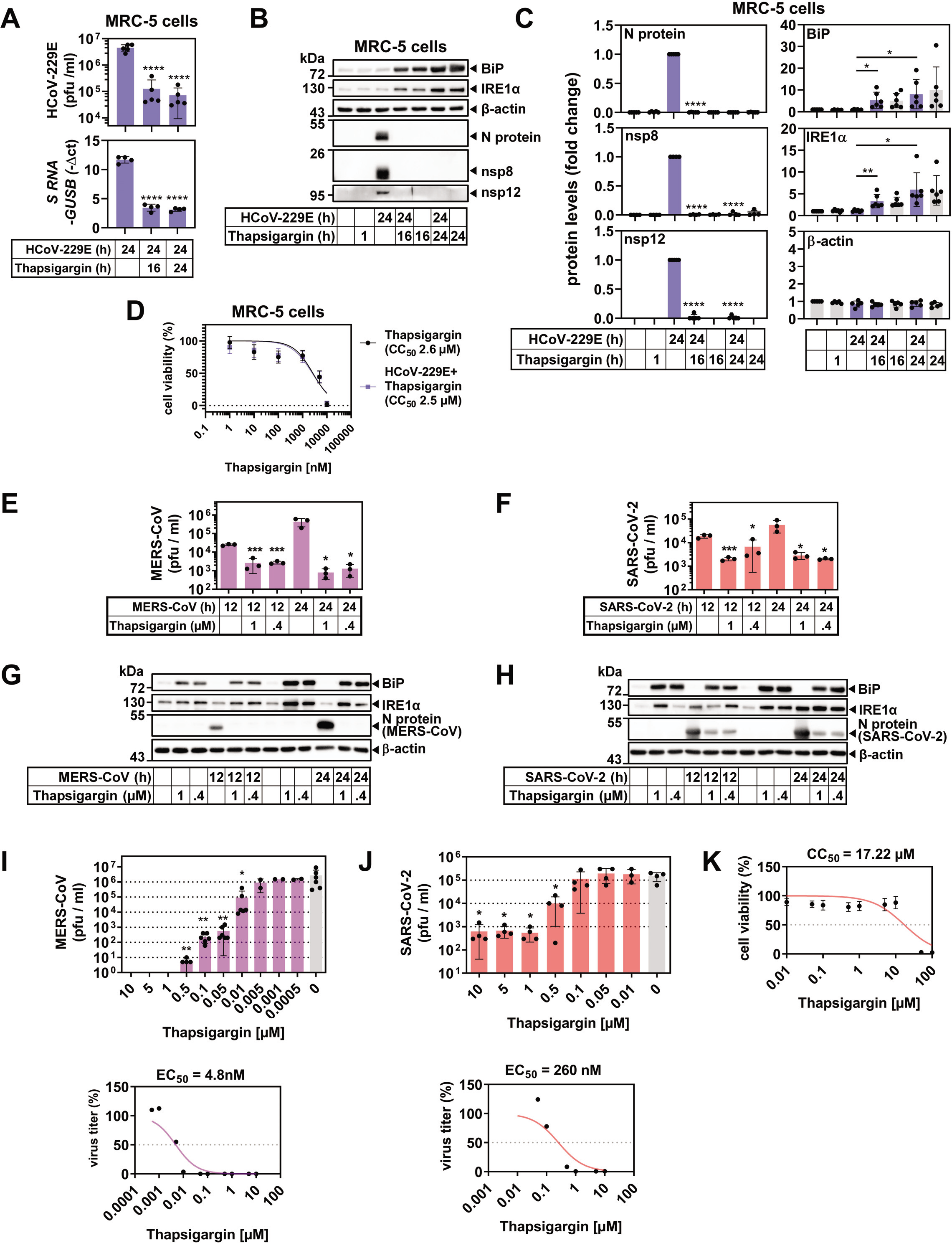
Thapsigargin inhibits the replication of high- and low-pathogenic human coronaviruses in multiple cell types. (A-D) Human embryonic MRC-5 lung fibroblasts were infected with HCoV-229E according to the scheme shown in Fig. 1B. Viral titers (A, upper graph), and expression of viral S gene-containing RNAs (A, lower graph), viral and host cell proteins (B, C) and cell viability (D) were analyzed and quantified as described in the legend of Fig. 1. (E-K) Similarly, HuH7 cells or Vero E6 African green monkey kidney epithelial cells were infected with MERS-CoV (MOI=0.5) or SARS-CoV-2 (MOI=0.5) for 12 h or 24 in the presence / absence of 0.4 μM or 1 μM thapsigargin. (E, F) show viral titers and (G, H) the corresponding expression of MERS-CoV / SARS-CoV-2 nucleocapsid (N) and host cell proteins, respectively. (I) Dose-dependent suppression of MERS-CoV-2 replication by thapsigargin (upper graph) and the estimated effective inhibitory concentration (EC_50_) in HuH7 cells infected with an MOI of 0.5 (lower graph). (J) Dose-dependent suppression of SARS-CoV-2 replication by thapsigargin (left graph) and the calculated effective inhibitory concentration (EC_50_) in Vero E6 cells infected with an MOI of 0.5 (right graph). (K) The CC_50_ of thapsigargin in Vero E6 cells was calculated by MTS assays as described in the legend of Fig. 2G-H. Data points show values from independent biological replicates, error bars show s.d.. Asterisks indicate p values (*p ≤ 0.05, **p ≤ 0.01, ***p ≤ 0.001, **** p ≤ 0.0001) obtained by two-tailed unpaired t-tests. See Fig. S2 for quantifications from replicates for MERS-CoV / SARS-CoV-2 immunoblot experiments.

To characterize the underlying molecular mechanisms responsible for the observed antiviral effects of thapsigargin, we focused on two highly pathogenic coronaviruses, MERS-CoV and SARS-CoV, for which, to our knowledge, no side-by-side comparison of proteomic changes has been reported to date. The large-scale proteomic study included (i) untreated cells and cells that were (ii) infected with MERS-CoV, (iii) SARS-CoV-2, (iv) treated with thapsigargin, (v and vi) infected with one of these viruses in the presence of thapsigargin. We used label-free quantification of six replicates per sample to determine the expression levels of > 5000 proteins from total cell extracts.

In a systematic approach, we identified differentially expressed proteins (DEPs) based on pairwise comparisons of proteins obtained from untreated cells, virus-infected cells or thapsigargin-treated cells using a p value of −log_10_ ≥ 1.3 as cut-off. As visualized by Volcano plot representations, MERS-CoV infection suppressed 413 (at 12 h p.i.) and 1171 proteins (at 24 h p.i.), respectively, and increased the levels of 150 proteins (at 12 h p.i.) and 508 proteins (at 24 h p.i.), respectively **(Fig. 4A, B)**, while SARS-CoV-2 suppressed the expression of 232 proteins at 12 h p.i. and 141 proteins at 24 h p.i. and increased the expression of 184 proteins at 12 h p.i. and 56 proteins at 24 h p.i. **(Fig. 4C, D)**. Thapsigargin treatment alone suppressed / induced large numbers of proteins in HuH7 cells (969 down, 923 up at 12 h; 1711 down, 958 up at 24 h) and in Vero E6 cells (201 down, 179 up at 12 h; 216 down, 149 up at 24 h) **(Fig. 4A-D)**. As expected, this analysis also identified viral proteins as the most strongly regulated DEPs. A comparison of virus-infected cells with virus-infected cells treated with thapsigargin revealed a complete suppression of all viral proteins and a large number of proteins with increased expression in thapsigargin-treated cells infected with MERS-CoV (676, 12 h p.i.; 1,100, 24 h p.i.; **blue groups of proteins**) or SARS-CoV-2 (268, 12 h; 326, 24 h; **blue groups of proteins**) **(Fig. 4A-D, right graphs)**. Also, similar numbers of proteins were identified with higher expression in virus-infected cells compared to virus-infected cells treated with thapsigargin **(Fig. 4A-D, right graphs; red groups of proteins)**. Together, these data lead us to conclude that thapsigargin causes a profound shift in protein expression in infected cells that likely contributes to the antiviral effects of this compound.

**Fig. 4.**
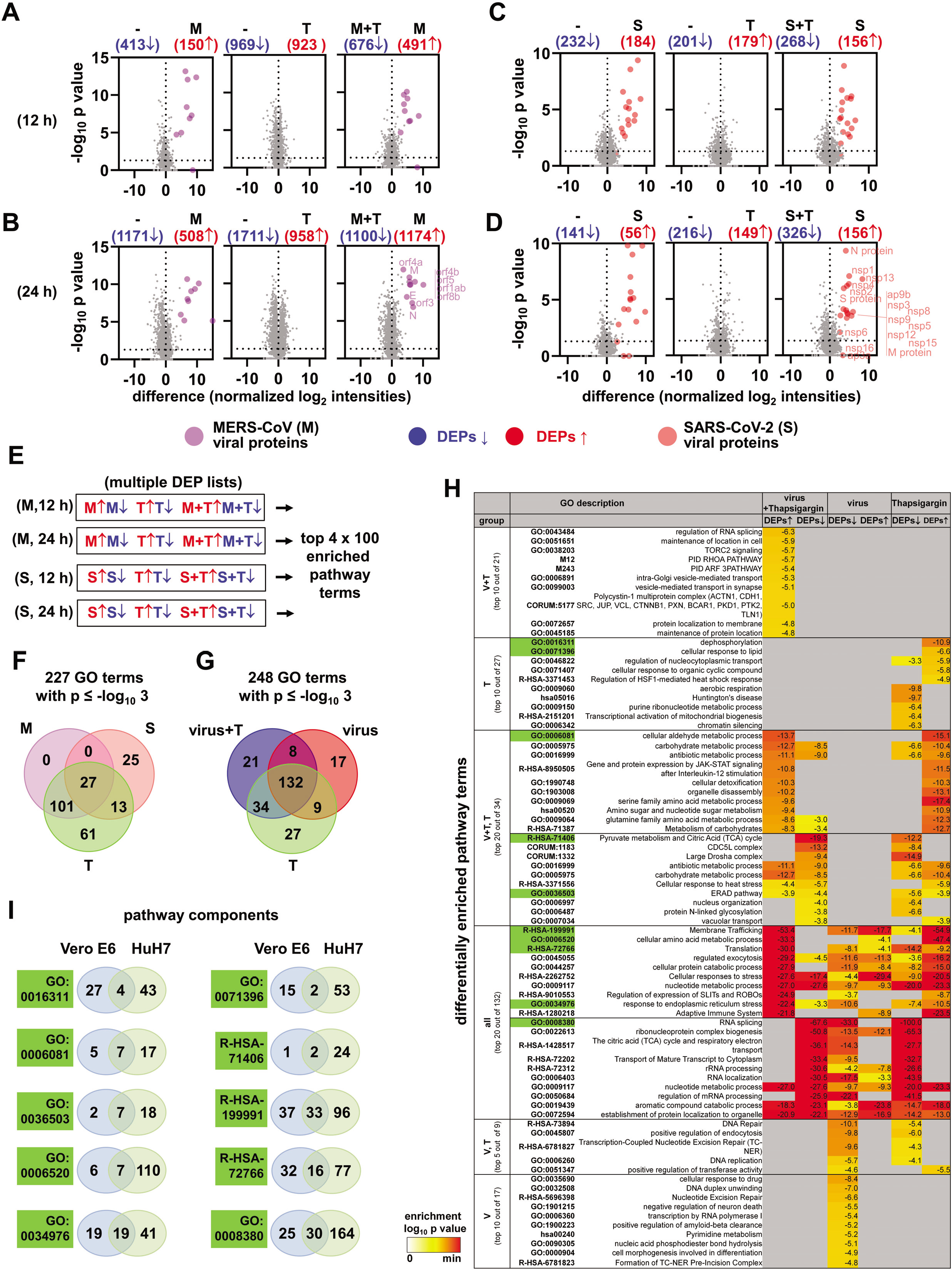
Proteome profiling of thapsigargin effects on MERS-CoV or SARS-CoV-2-infected cells reveals multiple virus- and thapsigargin-specific protein and pathway patterns. (A-D) Total cell extracts from uninfected cells (-), HuH7 cells infected with MERS-CoV (M, MOI=0.5) for 12 h (A) or 24 h (B), or Vero E6 cells infected with SARS-CoV-2 (S, MOI=0.5) for 12 h (C) or 24 h (D), in the presence or absence of thapsigargin (T, 1 μM) were subjected to LC-MS/MS analysis. In total, 5,372 (from HuH7) or 5,072 (from Vero E6 cells) majority protein IDs were identified. Quantile-normalized log_2_-tranformed protein intensities were used for all further calculations. Volcano plots show the distribution of pairwise ratio comparisons and the corresponding p values which were obtained by pooling data from two independent experiments and from three technical replicates per sample. Blue and red colors indicate differentially expressed proteins (DEPs, ratio > 0, p ≥ −log_10_ 1.3). Purple and light red colors indicate individual viral proteins. (E) Scheme of multiple gene ID lists (corresponding to the DEPs shown in (A)-(D)) that were used for overrepresentation analyses to identify the top 100 enriched pathway categories per virus and time point using Metascape software (Zhou *et al*., 2019). Complete lists of pathways are shown in Fig. S3 and S4 as clustered heatmaps. The top 5 enriched pathway categories for up- or downregulated DEPs are shown in Fig. S5A. (F, G) The 400 enriched pathway categories were pooled and filtered for common and distinct pathways considering only terms with enrichment p values ≤ log_10_ −3. (F) Venn diagram showing pathway terms specific to MERS-CoV (M), SARS-CoV-2 (S) or thapsigargin (T). The top 20 enriched pathway categories are shown in Fig. S5B. (G) Venn diagram showing pathway terms specific for virus, thapsigargin, or infection plus thapsigargin (virus + T) conditions. (H) The heatmap shows the top differentially enriched pathways corresponding to the Venn diagram shown in (G). Green colors refer to the pathways highlighted in (I). (I) Ten examples of differential and joint gene ID compositions of pathways enriched in HuH7 or Vero E6 cells.

We then devised a bioinformatics strategy to identify patterns of co-regulated or unique pathways and link deregulated protein sets identified in these data to specific (known) biological functions. As shown schematically in **Fig. 4E**, we sorted the DEPs from each of the four groups shown in **Fig. 4A-D** into four multiple gene ID lists, annotated the gene IDs to biological pathways and generated hierarchically clustered heatmaps of the top 100 differentially enriched pathway categories for the 12 h p.i. and 24 h p.i. time points of MERS-CoV- and SARS-CoV-2-infected cells, respectively, versus thapsigargin-treated cells by overrepresentation analysis (ORA) using Metascape software (Zhou *et al*, 2019). In this analysis, the groups of up- or downregulated proteins were kept separate to preserve information on whether specific DEPs belonging to particular overrepresented pathway terms were regulated in the same or opposite direction. Inspection of the four top 100 clustered heatmaps shows many similarities but also differences in pathways and their enrichment p values in response to virus infection or thapsigargin, all together demonstrating the complexity of the cellular response to CoV infections or chemical stressors **(Fig. S3–S4)**. By condensing this information to the top 5 pathways for up-/or downregulated DEPs we found that many of the most highly enriched categories are related to RNA, DNA, metabolic functions and localization **(Fig. S5A)**. We then combined the 400 pathway categories and searched this list for identical or unique GO terms in response to MERS-CoV, SARS-CoV-2 or thapsigargin. By filtering 227 pathways (out of 400) with enrichment p values ≤ −log_10_ 3 we found 27 pathway categories shared by both viruses and by thapsigargin, which are mostly related to RNA, folding, stress and localization **(Fig. 4F, Fig. S5B)**. 61 pathway categories unique to thapsigargin almost exclusively represented metabolic and biosynthetic pathways as shown for the top 20 overrepresented pathways containing up- or downregulated DEPs, suggesting that thapsigargin on its own, unlike CoV infection, initiates a broad metabolic response (**Fig. 4F, Fig. S5B)**.

This raised the question of whether the thapsigargin effects were retained in infected cells or, alternatively, drug-sensitive pathway patterns were reprogrammed (or masked) by the virus infection. To address this point, we pooled all pathways enriched under virus+thapsigargin conditions and compared them to virus infection or thapsigargin alone. 53% (132 out 248) pathway terms were shared by these three conditions reflecting multiple stress-related, catabolic and RNA regulatory processes **(Fig. 4G, H)**. 21 pathway terms were unique to the virus+thapsigargin situation. They primarily mapped to specific splicing, signaling (TORC, RHOA, ARF3) and transport/localization pathways **(Fig. 4G, H)**. The 34 categories shared by virus+thapsigargin and thapsigargin conditions but were not detectable in cells infected with virus (only) recapitulate the thapsigargin-regulated metabolic pathways (pyruvate, aldehyde, carbohydrate, amino/nucleotide sugar, amino acid and glutamine metabolism, TCA cycle, ERAD pathway, N-linked glycosylation) **(Fig. 4G, H)**. For several of these pathways (e.g. ERAD, heat stress, carbohydrate metabolism), some DEPs were induced while others were repressed, indicating remodeling of pathway functions at the protein level **(Fig. 4G, H)**. The 53 pathway terms that were absent in the virus+thapsigargin group of terms (groups 27, 9, 17 of the Venn diagram shown in **Fig. 4G**) represent a distinct set of terms, mostly related to nucleotide and DNA-related processes, such as DNA repair, DNA unwinding, chromatin silencing **(Fig. 4G, H)**. In summary, the functional analysis of DEPs at the level of differentially enriched pathway categories shows that the antiviral effects of thapsigargin strongly correlate with the activation / suppression of a range of metabolic programs.

The enriched pathway terms provided important overarching information on shared and unique biological processes but not necessarily encompassed identical sets of DEPs as exemplified by the ten pathways shown in **Fig. 4I**. We therefore refined our analysis to the individual component level to identify proteins with similar regulation between both viruses across both cell types. The proteomes of HuH7 and Vero E6 cells overlap by 57 % **(Fig. 5A)**. In this group, only 43 identical proteins were found to be deregulated by both MERS-CoV and SARS-CoV-2 **(Fig. 5B, left Venn diagrams)**. However, under thapsigargin+virus conditions, 108 proteins were upregulated and 61 proteins were downregulated, respectively **(Fig. 5B, right Venn diagrams)**. Using the example of the top 50 DEPs, it becomes apparent that the majority of proteins are regulated in the same direction by thapsigargin alone; demonstrating that thapsigargin largely overrides any virus-induced modulation of host processes **(Fig. 5C)**.

**Fig. 5.**
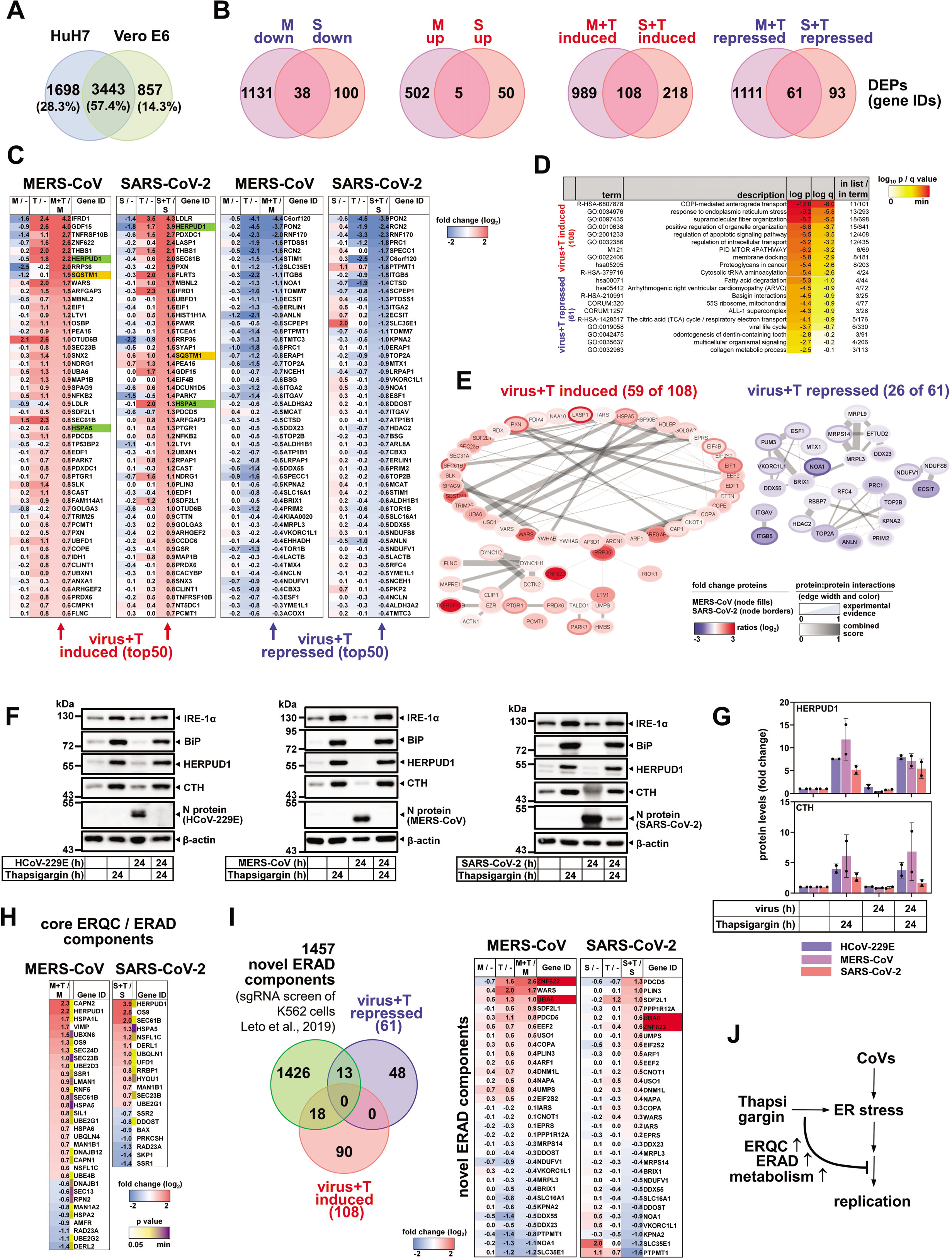
Thapsigargin effects on MERS-CoV or SARS-CoV-2-infected cells reveal the regulation of a specific network of proteins involved in transport, ERQC / ERAD and ER stress. (A) Venn diagram demonstrating the overlap of orthologues proteins expressed in HuH7 and Vero cells based on the NCBI gene IDs corresponding to majority protein IDs. (B) The overlap of virus- and thapsigargin-regulated proteins common to HuH7 and Vero E6 cells was calculated based on gene IDs. This analysis identifies 180 proteins with higher and 61 proteins with lower expression in thapsigargin-treated infected cells compared to virus infection alone (ratio > 0, p ≥ −log_10_ 1.3). (C) Heatmaps showing individual mean ratio values of log_2_-transfomed normalized protein intensities for the top 50 up- or downregulated proteins in virus-infected and thapsigargin-treated cells. Ratio values of infected or thapsigargin-treated conditions compared to untreated cells (-) cells are shown for comparison. Note that ratios are sorted and color coded according to the virus plus thapsigargin conditions with infection alone conditions set as denominator. Green colors highlight HERPUD1 and BiP (HSPA5) values. Orange colors highlight SQSMT1 which was also identified as SARS-CoV-2-regulated protein in an independent study (Stukalov *et al*., 2020). (D) Overrepresentation analysis showing the top 10 pathways mapping to gene IDs with increased (180 proteins, red) or decreased (61 proteins, blue) expression levels in thapsigargin-treated and infected cells compared to virus infection alone. Gene ID lists were analyzed using Metascape software (Zhou *et al*., 2019). (E) Experimental evidence, co-occurrence, co-expression and confidence scores from the STRING database (Szklarczyk *et al*, 2019) were used to identify protein:protein interactions (PPI) amongst the 180 up- and 61 downregulated thapsigargin-sensitive proteins. As shown, based on experimental evidence and combined STRING score criteria, 59 and 26 coregulated proteins are engaged in defined PPI networks; the remaining proteins are not known to interact. (F) Validation of thapsigargin-induced HERPUD1 and CTH upregulation by immunoblotting of HuH7 or Vero E6 whole cell extracts from cells treated as described above. BiP and IRE1α levels are shown for comparison. (G) Quantification of thapsigargin-mediated re-expression of HERPUD1 and CTH in cells infected with HCoV-229E, MERS-CoV or SARS-CoV-2 from two independent immunoblot experiments. Error bars show s.d.. (H) Heatmap showing thapsigargin-reprogrammed proteins of KEGG 04141 (mean ratio ≥ 1.5 fold) along with p values. See also Fig. S6 for projection of thapsigargin-mediated protein changes on the KEGG pathway map. (I) Venn diagram showing the intersection of thapsigargin- / virus-regulated proteins with all novel ERAD components (FDR of 1 %) identified by (Leto *et al*., 2019). The regulation of 31 overlapping components is shown as a heatmap displaying mean ratio values in thapsigargin-treated or infected cells. Red colors highlight UBA5 and ZNF622 as discussed in the text. (J) Summary of the main findings of our study. Abbreviations: M, MERS-CoV; S, SARS-CoV-2; T, thapsigargin.

In the absence of thapsigargin, the virus infection generally has little or opposite effects on the levels of the 108 proteins, as exemplified by the suppression seen for BiP (HSPA5) or HERPUD1 **(Fig. 5C, highlighted in green)**. The 108 induced factors map to pathways involving COPI-mediated anterograde transport, ER stress, organelle organization and apoptosis **(Fig. 5D)**. Across their pathway annotations, 59 out of the 108 proteins were reported to strongly interact, thus probably being involved in protein:protein networks that coordinate activities of the enriched pathways **(Fig. 5E, left graph)**. Likewise, the 61 repressed proteins map to specific (though different) pathways, such as fatty acid degradation or viral life cycle **(Fig. 5D)**. 26 components can be allocated to a few small protein interaction networks **(Fig. 5E, right graph)**.

We then validated mass spectrometry data by immunoblotting, confirming the induction of HERPUD1 in thapsigargin-treated cells infected individually with each of the three CoVs **(Fig. 5F, G)**. We also confirmed an additional hit belonging to the enriched pathways GO:0006520 (cellular amino acid metabolic process) and GO:0034976 (response to endoplasmic reticulum stress, as shown in **Fig. 4G, H**), cystathionine-γ-lyase (CTH), a regulator of glutathione homeostasis and cell survival (Lee *et al*, 2014), as a further independent example for the fidelity and robustness of the proteomics data **(Fig. 5F, G)**.

The highly inducible HERPUD1 protein has an essential scaffolding function for the organization of components of the core ER-associated protein degradation (ERAD) complex (Leitman *et al*, 2014; Okuda-Shimizu & Hendershot, 2007). ER quality control (ERQC) and ERAD pathways are critically involved in the qualitative and quantitative control of misfolded or excessively abundant proteins in the ER. If protein folding in the ER fails, the proteins are retro-translocated through a HERPUD1-dependent ER membrane complex to the cytosol for proteasomal degradation (Behnke *et al*, 2015). By searching our proteomics data for further ERAD factors we were able to retrieve a total of 34 (for MERS-CoV) and 20 (for SARS-CoV-2) proteins of the canonical ERQC and ERAD pathways for which a differential expression was observed in virus-infected cells treated with thapsigargin **(Fig. 5H)**. Mapping of these data on the KEGG 04141 pathway suggests that thapsigargin enhances or restores these mechanisms at key nodes of ERQC and ERAD in coronavirus-infected cells **(Fig. S6)**.

We also intersected the 108+59 proteins jointly regulated by thapsigargin in MERS-CoV and SARS-CoV-2 infected cells with data from a recent genome-wide sgRNA screen that reported new ERAD factors required for protein degradation (Leto *et al*, 2019). This analysis identified 31 additional thapsigargin-regulated factors that may further support antiviral ERAD, including UBA6 and ZNF622, which were recently described either as negative regulators of DNA virus infections or of autophagy, the latter process playing diverse roles during CoV infection **(Fig. 5I)** (Jia & Bonifacino, 2020; Maier & Britton, 2012; Mun & Punga, 2019).

In conclusion, these data show that thapsigargin forces the (re)expression of a dedicated network of proteins with roles in ER stress, ERQC, ERAD, and a range of metabolic pathways. Collectively, these changes at the protein level confer an “antiviral state” and profoundly suppress CoV replication as summarized schematically in **Fig. 5J**.

## Discussion

In this study, we report a potent inhibitory effect of the chemical thapsigargin on the replication of three human CoVs in three different cell types. Following up on observations that CoVs globally suppress UPR/ER stress factors, we find that thapsigargin counteracts the CoV-induced downregulation of BiP, HERPUD1 (and CTH) and increases IRE1α levels. In this context, thapsigargin also plays a role in overcoming the coronavirus-induced block of global protein biosynthesis. Proteome-wide data revealed a thapsigargin-mediated reprogramming of metabolic pathways and helped to identify a network of specific thapsigargin-regulated factors, including candidates from the ERQC/ERAD pathways that, most likely, are involved in the destruction of viral proteins. The positive effects of prolonged thapsigargin treatment on the expression of cellular BiP and HERPUD1 are well documented (Kokame *et al*, 2000; Li *et al*, 1993; Ma & Hendershot, 2004; Sun *et al*, 2015). Thus, the key finding of our study is that the thapsigargin-mediated induction of ER factors overrides suppressive effects of CoVs on ER functions, as illustrated here for BiP, IRE1α and HERPUD1, but also at the global proteomic scale.

BiP is one of the most abundant cellular proteins (also in our mass spectrometry data) and plays essential roles in development and disease (Wang *et al*, 2017; Zhu & Lee, 2015). In yeast, the inducible expression of the BiP homologue Karp2 is particularly essential for disposing of toxic proteins and reducing cellular stress (Hsu et al, 2012). Hence a reduction of BiP levels (as seen during CoV infection) and the contrary effect of thapsigargin-mediated upregulation are likely to have opposing consequences on cell fate upon infection. Similarly, IRE1α is suggested to mediate protective and adaptive responses suitable to alleviate ER stress, e.g. by balancing lipid bilayer stress, an aberrant perturbation of ER membrane structures, which may be expected to occur upon DMV formation in CoV-infected cells (Chen & Brandizzi, 2013; Halbleib *et al*, 2017; Snijder *et al*., 2020). Accordingly, high levels of BiP, HERPUD1 and IRE1α may increase in general the resilience of cells when infected by diverse pathogens. In line with this, our data show that, in cells infected with representative coronaviruses, a protective ER response is initially elicited at the mRNA level (Fig. 1C and (Poppe *et al*., 2017)). However, the global suppression at the protein level (or the lack of induction) indicates that CoVs have evolved strategies at the posttranscriptional or translational level to escape the protective antiviral activities of BiP, IRE1α and HERPUD1.

Together with PERK, all three proteins are key regulators of ERQC/ERAD pathways and ample evidence shows that their expression, regulation and activities are intimately linked (Hetz & Papa, 2018; Karagoz *et al*., 2019). A recent study reported that PERK activation induces the RPAP2 phosphatase, inactivates IRE1α kinase activity and aborts IRE1α-mediated adaptive functions in response to the chemical stressor Brefeldin A (Chang *et al*, 2018). In another report, high BiP levels exerted negative control of IRE1α by direct binding the kinase or by promoting IRE1α degradation (Chang *et al*., 2018; Sepulveda *et al*, 2018). Here, we show a different scenario, in which high BiP and IRE1α protein levels coincide with an antiviral state as well as with improved metabolic functions, suggesting unique modes of cross regulation of PERK, BiP and IRE1α in CoV-infected cells exposed to chemical stress.

The regulation of HERPUD1 and several ERAD factors by thapsigargin provides an additional layer of control contributing to the rapid suppression of CoV proteins. While ERAD is generally considered to dispose of unwanted proteins in the ER (Brodsky, 2012; Christianson *et al*, 2012), a process called “ERAD tuning” has been suggested to segregate ERAD components and thereby positively contribute to replicative organelle formation in RNA virus infections (Byun *et al*, 2014; Noack *et al*, 2014). Our data are compatible with a model in which a modulation of ERAD by small molecules may antagonize “ERAD tuning”, thereby preserving normal ERAD-mediated degradation.

Clearly, the mechanistic basis for these effects remains to be identified in additional studies. The proteomic data show that thapsigargin affects multiple pathways beyond the core ER stress response. They also indicate that it will not be trivial to identify the essential targets that mediate thapsigargin’s antiviral effects. Our data provide a rich resource for further drug target analysis, also in conjunction with the few deep protein sequencing studies available for SARS-CoV-2 (but not MERS-CoV) (Bojkova *et al*, 2020; Bouhaddou *et al*, 2020; Grenga *et al*, 2020; Stukalov *et al*, 2020). Our study fills an important knowledge gap by providing a direct side-by-side comparison of pharmacologically targeted cells infected with two highly pathogenic human coronaviruses.

In the absence of effective therapeutic and prophylactic strategies (antivirals and vaccines) to combat coronaviruses, and in view of the current SARS-CoV-2 pandemic, we report these observations to invite other laboratories to embark on a broader investigation of this potential therapeutic avenue. Given that thapsigargin concentrations in the lower nanomolar range were shown to abolish CoV replication in cultured cells, even when added later in infection (8 h p.i.), this work identifies thapsigargin as an interesting drug candidate. The Ca^2+^ mobilizing and cytotoxic features of plant-derived thapsigargin have been studied for 40 years (Andersen *et al*, 2015; Patkar *et al*, 1979). Several analogues have already been designed and efficient and scalable purification or synthesis is now available for application in humans (Chu *et al*, 2018; Chu *et al*, 2017; Lopez *et al*, 2018). Recently, a protease-cleavable prodrug of thapsigargin, mipsagargin, has been evaluated in phase I and II clinical trials for prostate cancer (Doan *et al*, 2015; Mahalingam *et al*, 2019; Mahalingam *et al*, 2016; Patkar *et al*., 1979). It is not uncommon to adapt anti-proliferative cytostatic drugs (e.g. azathioprine, cylophosphamid, methotrexate) for the treatment of autoimmune and inflammatory disorders by applying lower doses than those needed for treating cancer (Marder & McCune, 2007). Similarly, low doses of thapsigargin combined with short term systemic or topical application in the airways might reduce viral load early on or in critically ill patients with a favorable therapeutic index with respect to antiviral versus cytotoxic effects. CoV also activate inflammatory, NF-κB-dependent cytokine and chemokines at the mRNA level (Poppe *et al*., 2017), some of which (CXCL2, CCL20) escape translational shut-down and are secreted in a cell-type specific manner **(Fig. S7)**. Some of these cytokines may contribute to the cytokine storm observed in some COVID-19 patients (Mehta *et al*, 2020). While thapsigargin had no effect on IL-8, IL-6, CXCL2 and CCL20 in cell culture **(Fig. S7)**, a single bolus of the compound was shown to efficiently reduce the translation of pro-inflammatory cytokines in preclinical models of sepsis (Wei *et al*, 2019). Thus, an additional benefit of thapsigargin treatment may arise from dampening overshooting tissue inflammation in COVID-19 patients. In summary, the study provides several lines of evidence that thapsigargin hits a central mechanism of CoV replication, which may be exploited to develop novel therapeutic strategies. This compound or derivatives with improved specificity, pharmacokinetics and safety profiles may also turn out to be valuable to mitigate the consequences of potential future CoV epidemics more effectively.

## Author Contributions

MS-S, CM, CM-B, HW, BV-A, NK performed experiments, analyzed and visualized data. UL performed mass spectrometry measurements and together with AW and IB analyzed proteomics data sets. NH analyzed RNA-seq data. TH sequenced SARS-CoV-2. MK conceived and supervised the study and analyzed and visualized the data. MK wrote the initial manuscript draft. JZ, MLS (and all other authors) helped to finalize the manuscript. All authors approved the submitted version of the manuscript.

## Acknowledgments

This work was supported by the following grants from the Deutsche Forschungsgemeinschaft (DFG, German Research Foundation): KR1143/9-2 and ZI618/6-2 (KFO309, P3 (to M.K. and J.Z.), Z1 (to T.H.); project 284237345), TRR81/3 (A07 (to M.L.S.), B02 (to M.K.), project 109546710); SFB1213/2 (B03 (to M.K., M.L.S.), project 268555672); SFB1021/2 (A01 (to J.Z.), C01 (to M.L.S.), C02 (to M.K.), Z02 (to T.H.), Z03 (to M.K., U.L.), project 197785619); GRK 2573 (RP4 (to M.L.S), RP5 (to M.K.), project 416910386) and INST 160/708-1 FUGB (to U.L.). The work was also supported by the German Ministry for Education and Research (RAPID, COVINET, and DZIF TTU 01.806, to J. Z.) and the LOEWE program of the state of Hesse (DRUID, B02, to J.Z.). Work in the laboratories of M.K. and M.L.S. is also supported by the IMPRS program of the Max Planck Society and the Excellence Cluster Cardio-Pulmonary Institute (EXC 2026: Cardio-Pulmonary Institute (CPI), project 390649896) and the DZL/UGMLC program. Work of J.Z., M.K. and S.B. is further supported by the Von-Behring-Roentgen-Stiftung.

We thank Andrea Nist and Thorsten Stiewe of the Genomics Core Facility, Philipps University Marburg, Marburg, Germany, for RNA sequencing and Christian Drosten for providing SARS-CoV-2 and MERS-CoV.

## Declaration of Interests

The authors declare no competing interests.

---

**Fig. S1.**
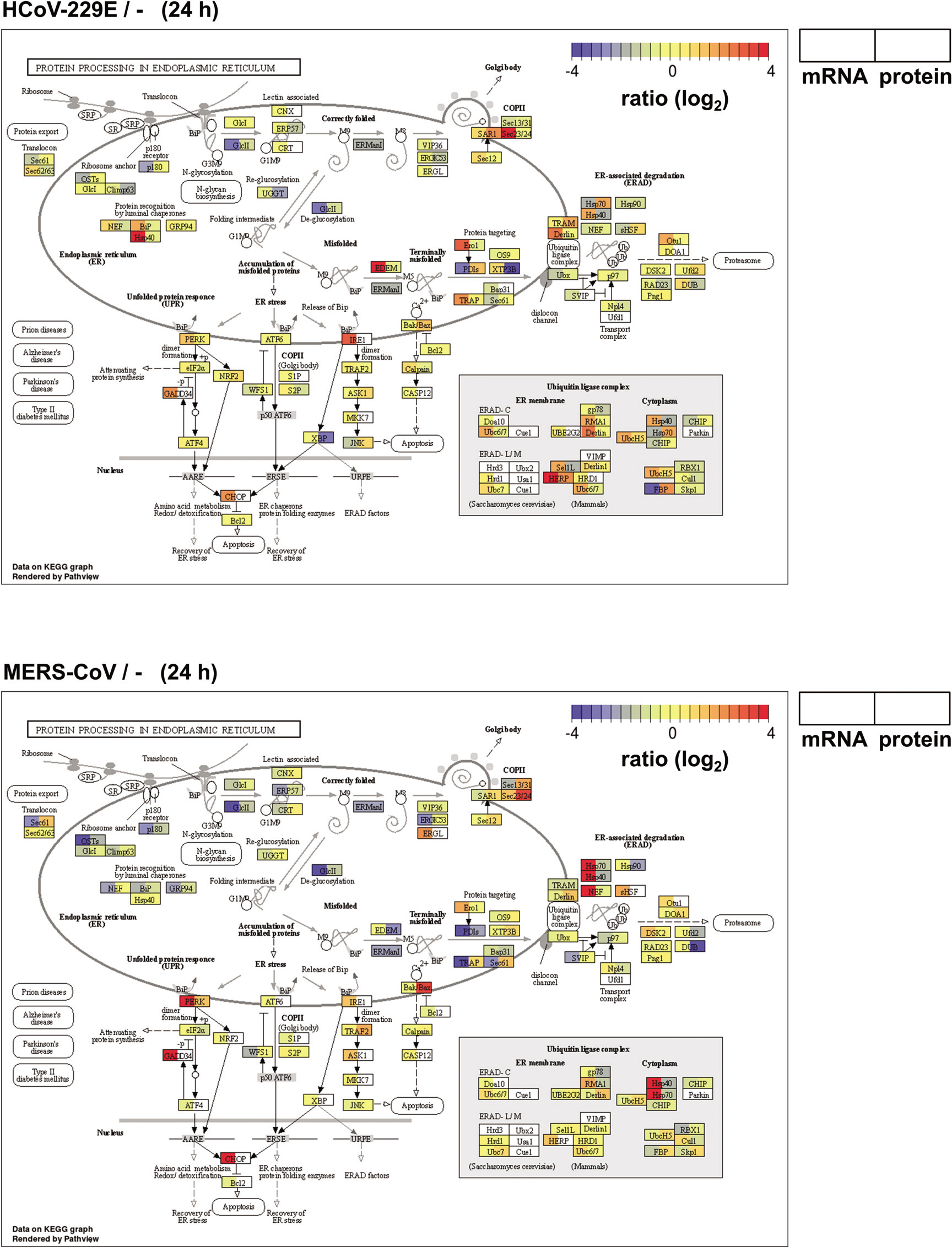
Opposing effects of HCoV-229E or MERS-CoV on ER stress genes at the mRNA and protein level. Projection of normalized ratio values for transcriptomic (by RNA-seq) and proteomic (by LC-MS/MS) data derived in parallel from HuH7 cells infected for 24 h with HCoV-229E or MERS-CoV with a MOI=1 on the components of the KEGG pathway 04141 “protein processing in endoplasmic reticulum”. The left side of the boxes show mRNA values, right sides show protein values.

**Fig. S2.**
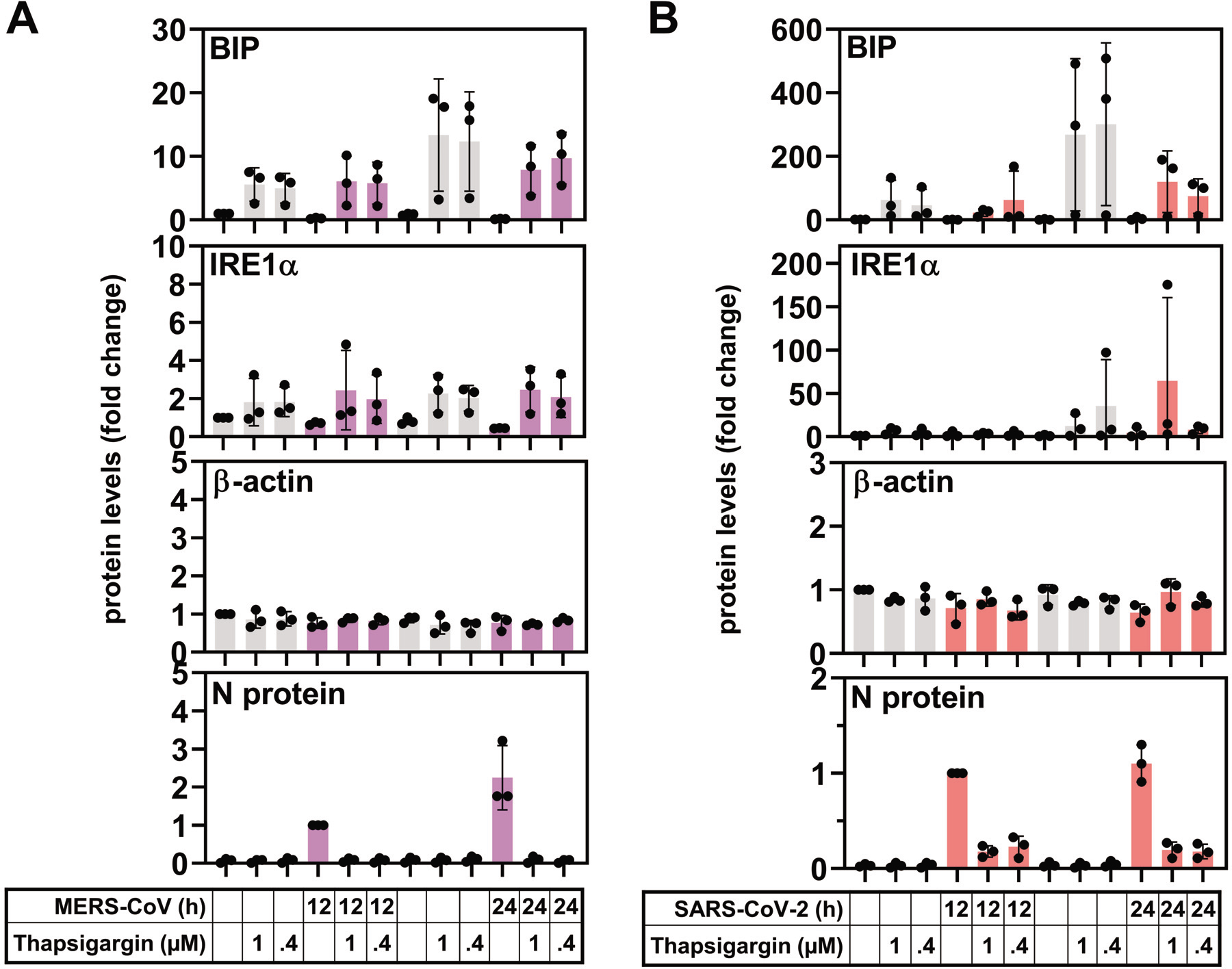
Thapsigargin suppresses MERS-CoV and SARS-CoV-2 N protein and upregulates BiP in infected cells. (A, B) show the quantification of replicate immunoblot experiments performed as shown in Fig. 3G and 3H. Data points show values from independent biological replicates, error bars show s.d..

**Fig. S3.**
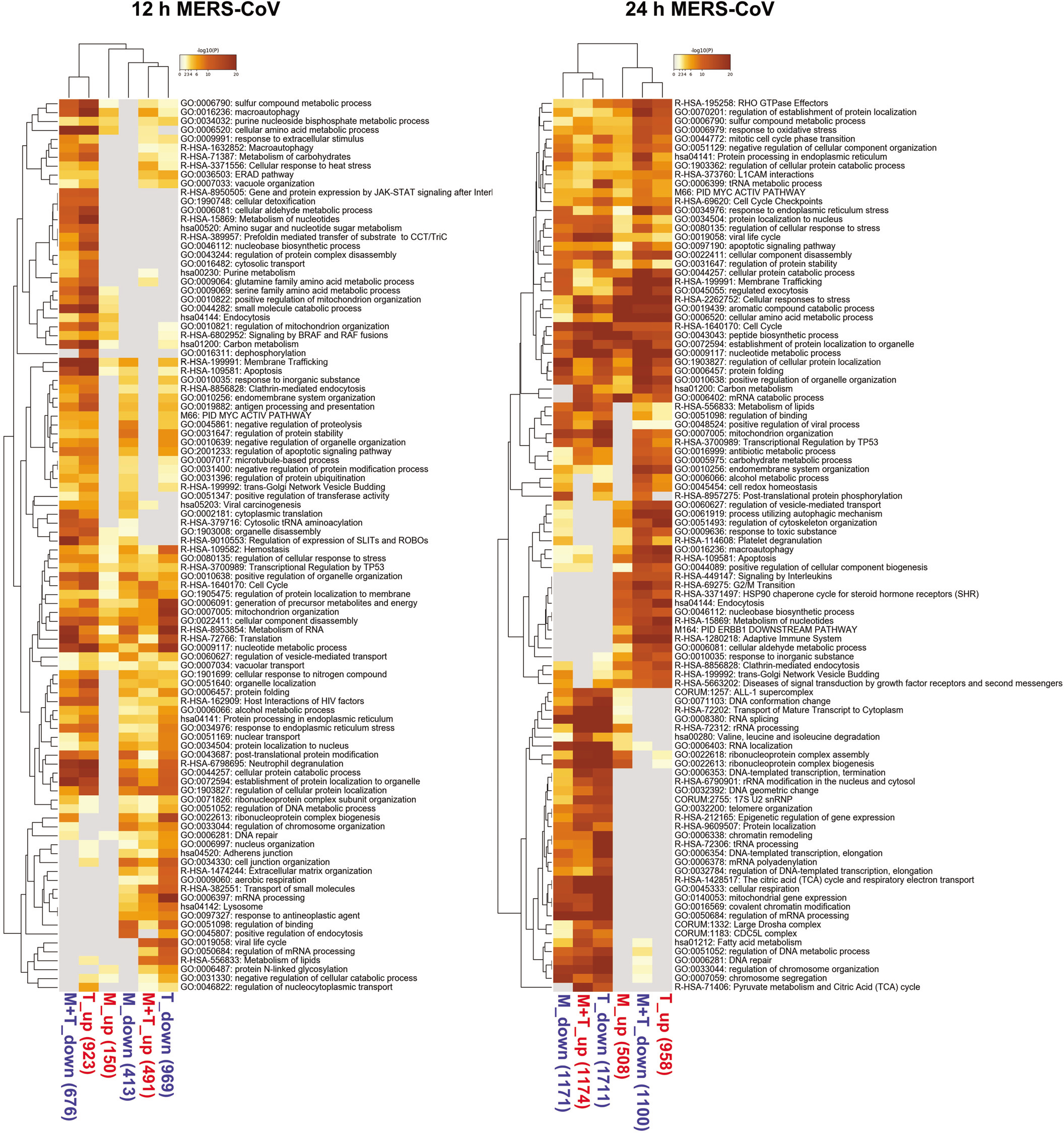
Identification of deregulated cellular pathways in MERS-CoV-infected HuH7 cells. Top 100 overrepresented pathways containing up- or downregulated DEPs (ratio > 0, p ≥ log_10_ 1.3) for the 12 h p.i. and 24 h p.i. time points of MERS-CoV-infected cells based on gene IDs derived from protein IDs. Blue and red colors indicate differentially expressed proteins as shown in Fig. 4A-B.

**Fig. S4.**
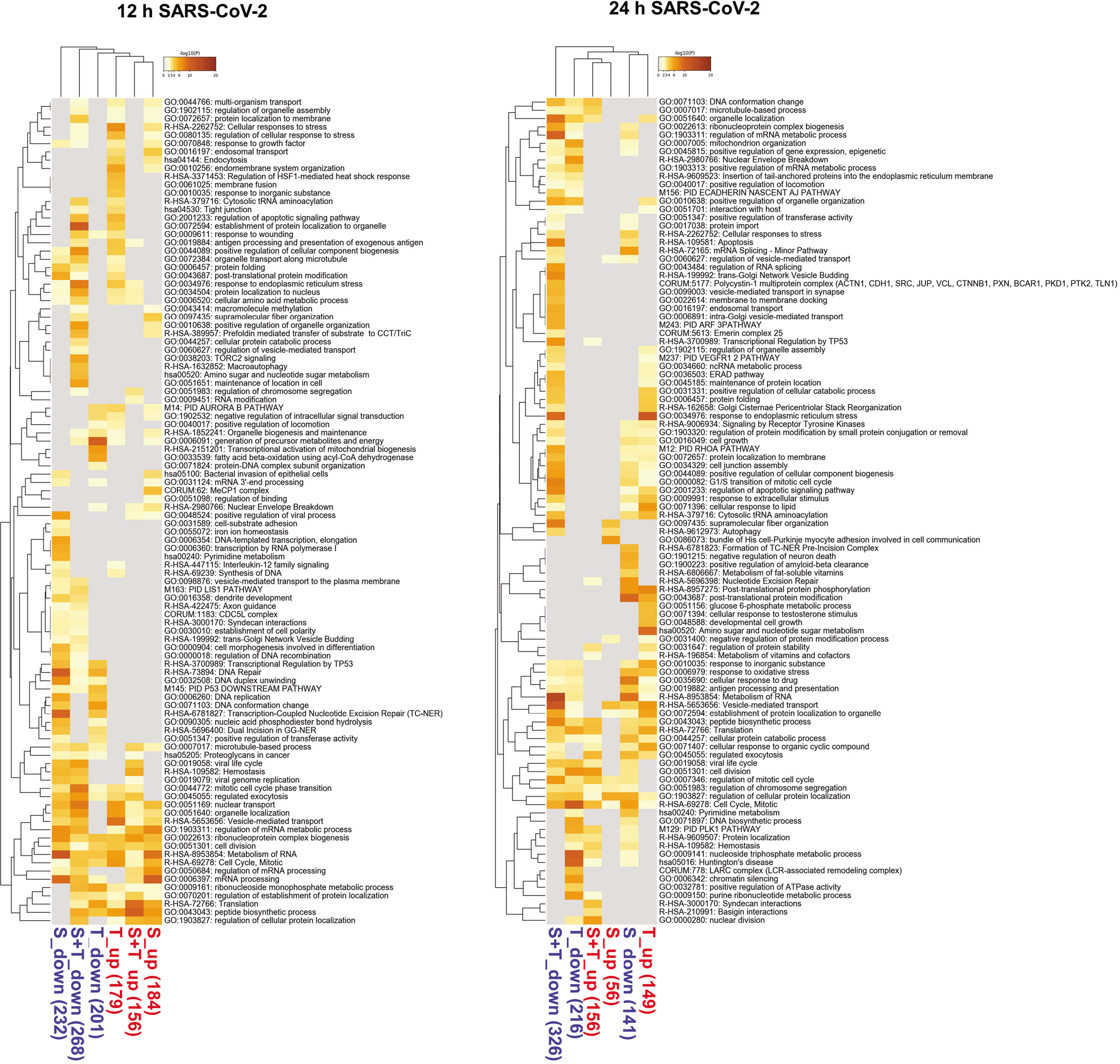
Identification of deregulated cellular pathways in SARS-CoV-2-infected Vero E6 cells. Top 100 overrepresented pathways containing up- or downregulated DEPs (ratio > 0, p ≥ log_10_ 1.3) for the 12 h p.i. and 24 h p.i. time points of SARS-CoV-2-CoV-infected cells based on gene IDs derived from protein IDs. Blue and red colors indicate differentially expressed proteins as shown in Fig. 4C-D.

**Fig. S5.**
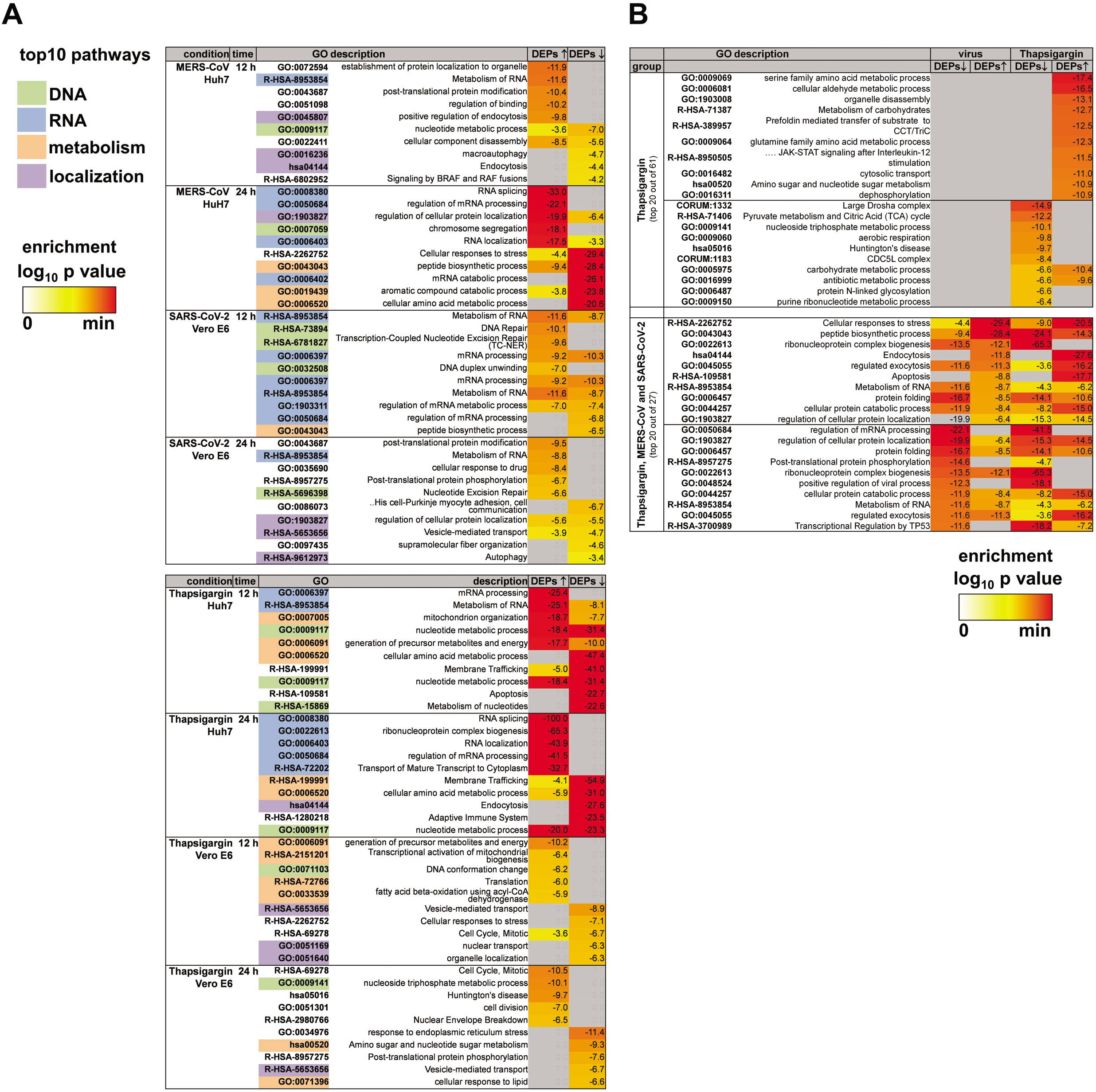
Top pathways regulated by MERS-CoV, SARS-CoV-2 or thapsigargin. (A) Top ten enriched pathways containing up- or downregulated DEPs extracted from the 100 enriched deregulated pathways shown in Fig. S3 / Fig. S4. Colors indicate highly common categories. (B) Top 20 pathways enriched with thapsigargin alone or jointly by MERS-CoV, SARS-CoV-2 and thapsigargin according to the Venn diagram shown in Fig. 4F.

**Fig. S6.**
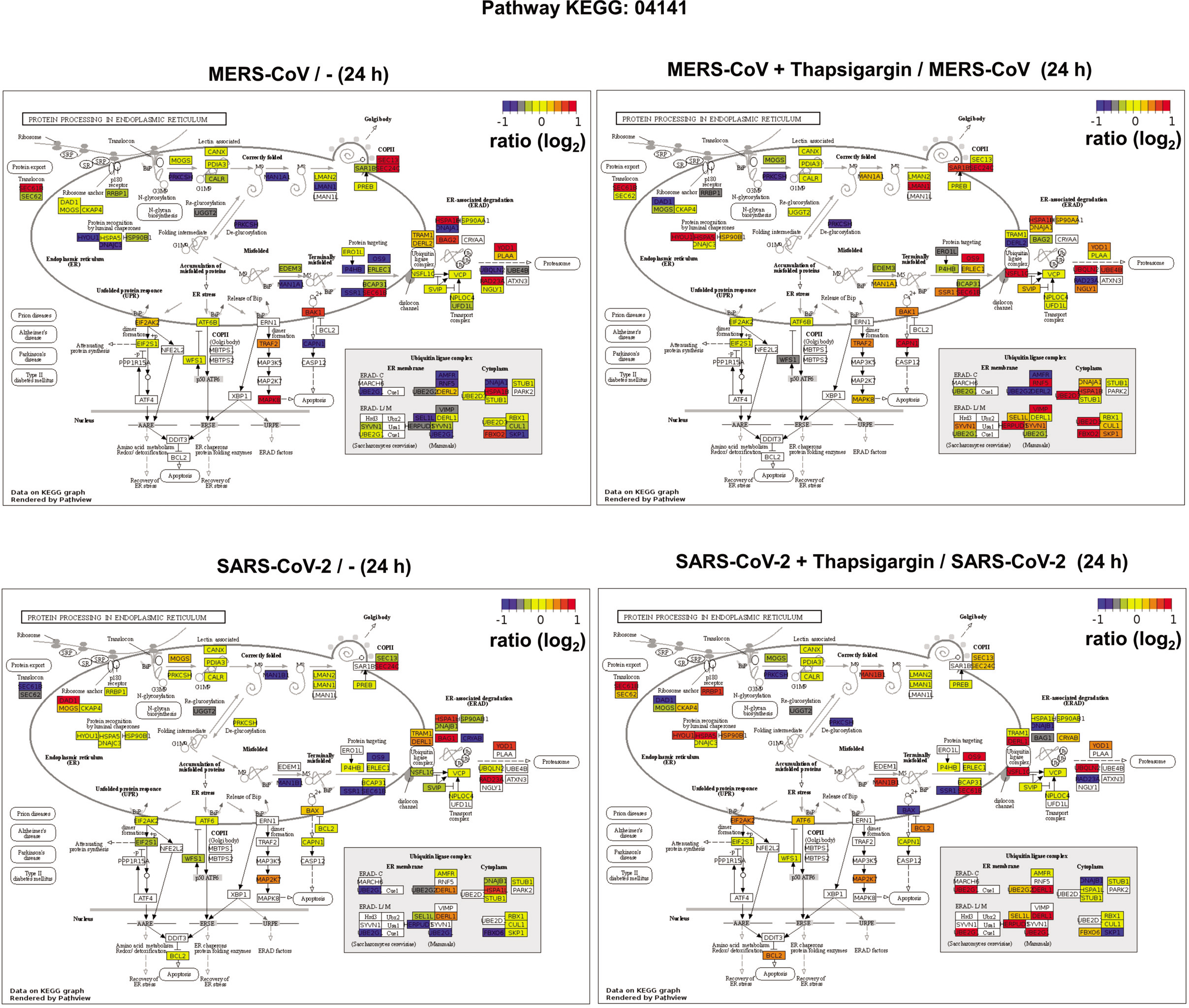
Projection of thapsigargin effects on protein levels of pathway KEGG 04141. Mean ratio values of all pathway components measured by LC-MS/MS in untreated cells and 24 h p.i. were projected on the KEGG 04141 pathway map (left graphs). The right graphs show the corresponding changes imposed by thapsigargin treatment of infected cells.

**Fig. S7.**
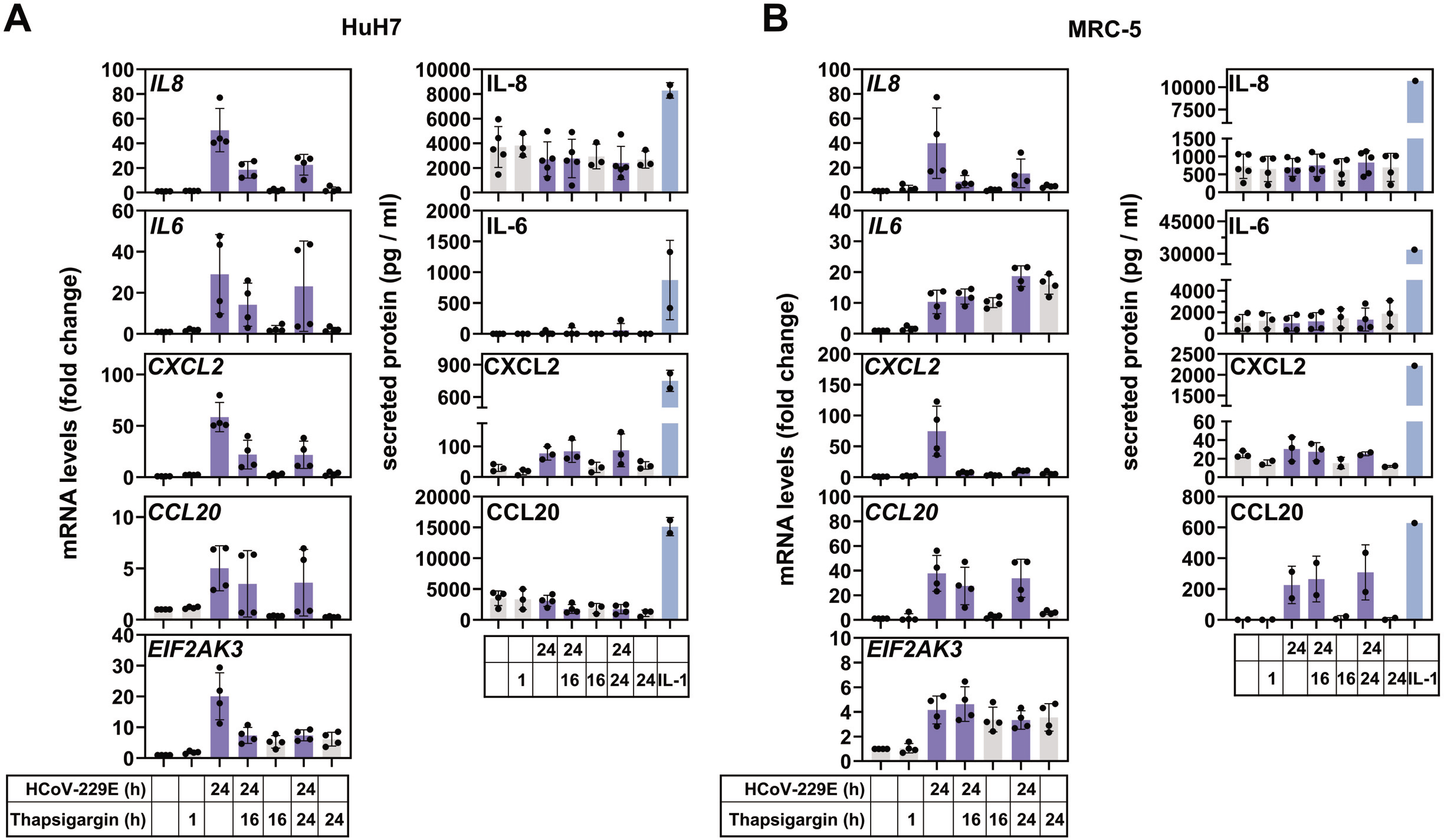
Analysis of inflammatory host cell transcripts showing thapsigargin-independent uncoupling of mRNA and protein levels in HCoV-229E-infected cells. HuH7 (A) or MRC-5 (B) cells were infected as described in Fig. 2B. Total RNA was used to analyze expression of inflammatory transcripts by RT-qPCR (left graphs) and the secretion of the corresponding proteins by ELISA (right graphs). Additionally, expression of the *EIF2AK3* mRNA encoding the PERK protein kinase was determined. PERK protein levels of HuH7 cells are shown Fig. 2E-F. Data points show values from independent biological replicates, error bars show s.d..

## Materials and Methods

### Cells and viruses

HuH7 human hepatoma cells (Japanese Collection of Research Bioresources (JCRB) cell bank; (Nakabayashi *et al*, 1982)) were maintained in Dulbecco’s modified Eagle’s medium (DMEM, including 3.7 g / l NaHCO_3_; PAN Biotech Cat No P04-03550) complemented with 10% filtrated bovine serum (FBS Good Forte; PAN Biotech, Cat No. P40-47500), 2 mM L-glutamine (Gibco, 21935-028), 100 U / ml penicillin and 100 μg /ml streptomycin. MRC-5 human embryonic lung fibroblasts ((ATCC, CCL-171)) were maintained in DMEM containing 1.5 g / l (w / v) NaHCO_3_ and complemented with 10% fetal calf serum (FCS; PAN Biotech Cat No. 1502-P110704), 2 mM L-glutamine, 100 U / ml penicillin and 100 μg / ml streptomycin, 1% minimum essential medium nonessential amino acids (100x MEM NEAA; Gibco Cat No 11140-035) and 1 mM sodium pyruvate (100 mM; Gibco 11360-039). Vero E6 African green monkey kidney epithelial cells (ATCC CRL-1586) were grown in DMEM, 10% FBS, 100 U / ml penicillin and 100 μg /ml streptomycin. HuH7 and MRC-5 cells were confirmed to be free of mycoplasma using the Venor^®^ GeM Classic kit (Minerva Biolabs).

Genome sequences of coronavirus strains used in this study are as follows: HCoV-229E (NCBI accession number AF304460.1, NCBI reference sequence NC_002645.1), MERS-CoV (NCBI accession number JX869059, NCBI reference sequence NC_019843.3). SARS-CoV-2 genome sequencing data have been submitted to the NCBI Short Read Archive repository under bioproject PRJNA658242 (SRA accession number SRP278165). MERS-CoV and SARS-CoV-2 were kindly provided by Christian Drosten.

### Virus infections and assessments of antiviral activity

To analyze the antiviral activity of thapsigargin, HuH7 cells (for HCoV-229E, MERS-CoV), MRC-5 cells (for HCoV-229E) and Vero E6 (for SARS-CoV-2) were infected at the indicated multiplicities of infection (MOI) and incubated at 33°C (for HCoV-229E and SARS-CoV-2) or 37°C (for MERS-CoV) in the presence or absence of thapsigargin, or with the appropriate volume of solvent control (DMSO) as indicated. At 24 h p.i., supernatants were collected and stored at −80 °C. Virus titers in the supernatants were determined by plaque assay. HCoV-229E was titrated on HuH7 cells seeded on 12-well plates. MERS-CoV was titrated on HuH7 cells seeded on 24-well plates, and SARS-CoV-2 was titrated on Vero E6 cells seeded on 24-well plates using standard procedures. Briefly, confluent monolayers of the appropriate cells were incubated with serial dilutions of virus-containing supernatants (diluted 10^1^ to 10^7^) and incubated at 33°C (HCoV-229E, SARS-CoV-2) or 37°C (MERS-CoV). After 1 h, the virus inoculum was replaced with fresh medium (MEM, Gibco) containing 1.25% Avicel^®^ (FMC Biopolymer, #RC591), 100 U/ml penicillin, 100 μg/ml streptomycin, and 10% FBS or FCS. At 48 h p.i. (for MERS-CoV) or 72 h p.i. (for SARS-CoV-2 and HCoV-229E), cell culture supernatants were removed. Cells were washed with PBS and fixed in freshly prepared 3.7% PFA in PBS overnight. Next, the fixing solution was removed, the cell layer was washed with PBS and stained with 0.15% (w/v) crystal violet (diluted in 20% Ethanol) and plaques were counted. For EC_50_ calculation, virus titers determined for virus-infected cells treated with DMSO only (no inhibitor) were used for normalization. EC_50_ values were calculated by non-linear regression analysis using GraphPad Prism 5.0 or 8.4.3 (GraphPad Software). All virus work was performed in biosafety level 2 (BSL-2; HCoV-229E) or biosafety level 3 (BSL-3; MERS-CoV, SARS-CoV-2) containment laboratories approved for such use by the local authorities (RP Giessen, Germany).

### Materials

Thapsigargin (Cayman Chemicals; Cay10522-1) was dissolved in DMSO as a 10 mM stock solution. Appropriate DMSO concentrations served as vehicle controls in some experiments. The following inhibitors were used: leupeptin hemisulfate (Carl Roth, #CN33.2), microcystin (Enzo Life Sciences, #ALX-350012-M001), pepstatin A (Applichem, #A2205), PMSF (SigmaAldrich, #P-7626). Pepstatin A, PMSF and microcystin were dissolved in ethanol and leupeptin in water. Other reagents were from Sigma-Aldrich or Thermo Fisher Scientific, Santa Cruz Biotechnology, Jackson ImmunoResearch, or InvivoGen and were of analytical grade or better.

Primary antibodies against the following proteins or peptides were used: anti β-actin (Santa Cruz, #sc-4778), anti PERK (Santa Cruz, #sc-377400), anti PERK (Abcam, #ab65142), anti BiP (Cell Signaling, #3177), anti eIF2α (Cell Signaling #9722), anti P(S51)-eIF2α (Cell Signaling #9721), anti P(S724)-IRE1α (Novus Biologicals, #NB100-2323), anti IRE1α (Santa Cruz, #sc-390960), anti ATF4 (Santa Cruz, #sc-390063), anti ATF3 (Santa Cruz, #sc-188), anti HERPUD1 antibody (Abnova, #H00009709-A01), anti CTH antibody (Cruz, #sc-374249), anti HCoV-229E N protein ((Ingenasa, Batch 250609), mouse anti HCoV-229E nsp12 (gift from Carsten Grötzinger), rabbit anti HCoV-229E nsp8 (Ziebuhr & Siddell, 1999), anti MERS-CoV N protein (Sinobiological, #100213-RP02), rabbit anti SARS-CoV N protein cross-reacting with SARS-CoV-2 N protein (gift from Friedemann Weber), anti SARS-CoV-2 N protein (Rockland, #200-401-A50), anti puromycin (Kerafast Inc., 3RH11, #EQ 0001), anti J2 (SCICONS, English & Scientific Consulting Kft, #10010200).

The following secondary antibodies were used: Dako P0447; polyclonal goat anti-mouse immunoglobulins/HRP, Dako P0448; polyclonal goat anti-rabbit immunoglobulins/HRP, Cy3-coupled anti rabbit (rb) IgG (dk, Merck Millipore, #AP182C), Dylight 488-coupled anti mouse (ms) IgG (dk, ImmunoReagent, #DkxMu-003D488NHSX),

### Cell lysis, *in vivo* puromycinylation, and immunoblotting

For whole cell extracts, samples derived from experiments performed with HCoV-229E were lysed in Triton cell lysis buffer (10 mM Tris,pH 7.05, 30 mM NaPPi, 50 mM NaCl, 1% Triton X-100, 2 mM Na3VO4, 50 mM NaF, 20 mM ß-glycerophosphate and freshly added 0.5 mM PMSF, 2.5μg/ml leupeptin, 1.0 μg/ml pepstatin, 1 μM microcystin). After 10-15 min on ice, lysates were cleared by centrifugation at 15,000 x g for 15 min at 4°C. Protein concentrations of supernatants were determined by Bradford assay and samples stored at −80 °C.

To label nascent polypeptides in intact cells (Iwasaki & Ingolia, 2017), HuH7 cells were seeded in 6 cm cell culture dishes (3 × 10^5^ cells) and treated as described in the figure legends. Thirty minutes prior to harvest, the medium was supplemented with 3 μM puromycin (InvivoGen, #ant-pr-1). Then, cells were lysed as described above. After immunoblotting (see below), membranes were stained with Coomassie brilliant blue and then hybridized with an anti puromycin antibody (Kerafast, #EQ0001) to detect puromycinylated polypetides.

Total cell lysates of MERS-CoV- and SARS-CoV-2-infected cells used for immunoblotting or mass spectrometry were prepared as follows. Cells were scraped in ice-cold PBS and pelleted at 500 x g for 5 min at 4°. Cell pellets were washed in ice-cold PBS and stored in liquid N2 (or lysed and processed immediately). After thawing, cell pellets (corresponding to ≈ 300.000 cells seeded in 60 mm dishes at the start of the experiment) were resuspended in 90 μl of ice-cold Ca^2+^ / Mg^2+^-free PBS and transferred to fresh tubes. After addition of 10 μl of 10% SDS, samples were heated at 100 °C for 10 min and centrifuged at 600 x g for 1 minute at room temperature. Supernatants were transferred to a fresh tube and heated again at 100°C for 10 minutes and centrifuged at 600 x g for 1 minute at room temperature. Protein concentrations were determined with the detergent compatible Bradford assay kit (Pierce™, #23246) using a 150-fold dilution. Aliquots corresponding to 20-25 μg protein (per lane) were mixed with 4 x SDS sample buffer (ROTI^®^Load, Roth, #K929) and stored at −20 °C prior to SDS-PAGE, or loaded immediately. Cell lysates were subjected to SDS-PAGE on 8-10% gels. The PageRuler™ prestained protein ladder (Thermo Scientific, #26616) was used as Mr marker.

Immunoblotting was performed essentially as described (Hoffmann *et al*, 2005). Proteins were separated on SDS-PAGE and electrophoretically transferred to PVDF membranes (Roti-PVDF, #T830 Roth). Membranes were stained with 0,1 % (w/v) Ponceau S (Sigma) dissolved in 5% acetic acid or Coomassie brilliant blue to confirm transfer and equal loading of proteins. After blocking with 5% dried milk in Tris-HCl-buffered saline/0.05% Tween (TBST) for 1 h, membranes were incubated for 12-24 h with primary antibodies, washed in TBST and incubated for 1-2 h with the peroxidase-coupled secondary antibody. Proteins were detected by using enhanced chemiluminescence (ECL) systems from Millipore or GE Healthcare. Images were acquired with the ChemiDoc TouchImaging System (BioRad) and quantified using the software ImageLab (versions V_5.2.1 or V_6.0.1, Bio-Rad).

### mRNA expression analysis by RT-qPCR

0.5 - 1 μg of total RNA was prepared by column purification (MachereyNagel) and transcribed into cDNA using Moloney murine leukemia virus reverse transcriptase (RevertAid Reverse Transcriptase, Thermo Fisher Scientific, #EP0441) in a total volume of 10 or 20 μl. 1 or 2 μl of this reaction mixture was used to amplify cDNAs using Taqman assays on demand (0.25 or 0.5 μl) (Applied Biosystems/Thermo Fisher Scientific) for *GUSB* (81 bp, Hs99999908_m1), *IL6* (95 bp, Hs00174131_m1), *IL8* (101 bp, Hs0017 4103_m1), *CXCL2* (68 bp, Hs00236966_m1), and *CCL20* (81 bp, Hs00171125_m1), as well as TaqMan Fast Universal PCR Master Mix (Applied Biosystems/ Thermo Fisher Scientific). Alternatively, primer pairs were designed and used with 2 μl of cDNA and Fast SYBR Green PCR Master Mix (Applied Biosystems/Thermo Fisher Scientific) to detect the mRNA of *EIF2AK3* (encoding PERK) (fw5’-AGAGATTGAGACTGCGTGGC-3’, re 5’-TCCCAAATACCTCTGGTTTGCT-3’) and of HCoV-229E S RNA (fw5’-TTTCAGGTGATGCTCACATACC-3’, re 5’-ACAAACTCACGAACTGTCTTAGG-3’). All PCRs were performed in duplicate on an ABI 7500 real-time PCR instrument. The cycle threshold value (ct) for each individual PCR product was calculated by the instrument’s software, and the ct values obtained for inflammatory/target mRNAs were normalized by subtracting the ct values obtained for GUSB. The resulting Δct values were also used to calculate relative changes of mRNA expression as ratio (R) of mRNA expression of treated / untreated cells according to the following equation: R=2^-((Δct treated)-(Δct untreated))^.

### Immunofluorescence

Cells were seeded for in μ-slides VI (Ibidi) and pre-cultured at 37°C, 6 % CO_2_. Virus infection as well as simultaneous thapsigargin treatment (1 μM) was performed for 24 h at 33°C, 6% CO_2_. After 2x washing, cells were fixed with 4% paraformaldehyde in PBS (Santa Cruz, #281692) for 5 min, washed 3x 10 min with Hank’s BSS (PAN, #P04-32505), blocked with 10% normal donkey serum (Jackson ImmunoResearch, #017-000-121) for 20 min and incubated with primary and secondary antibodies diluted in Hank’s BSS containing 0.005% saponin (Sigma-Aldrich, #S4521-10G) for 2 h at room temperature. Following 3 washing steps with Hank’s BSS containing 0.005% saponin, Cy3-conjugated (Millipore, #AP182C, 1:100) and Dylight488-conjugated (ImmunoReagents #DkxMu-003D488NHSX, 1:100) secondary antibodies were used. For controls, primary antibodies were omitted. Nuclei were stained with Hoechst 33342 (Invitrogen). Immunofluorescence images were analyzed using Leica DMi8 and the Leica LASX software. For double immunofluorescence analyses appropriate filter cubes were used (Dylight488: excitation 480/40 and emission 527/30, Cy3: excitation 560/40 and emission 630/75, as well as counterstaining with Hoechst 33342: excitation 405/60 and emission 470/50).

### Cell viability assays

MTS assays of HCoV-229E experiments were performed using the The CellTiter 96^®^ AQueous One Solution Cell Proliferation Assay kit (Promega, #G3582). In brief, 1.2 x 10^4^ HuH7 or 1 x 10^4^ MRC-5 cells were seeded in 96-well plates for 24 hours and thereafter treated with DMSO, thapsigargin, virus alone or virus plus thapsigargin for 24 hours as indicated in the figure legends. Then, the medium was replaced by 100 μl complete cell culture medium including 20 μl CellTiter 96^®^ AQueous one solution reagent according to the manufacturer’s recommendations. Cells were further incubated for 0.5 - 1 hour at 33° C. Then, absorbance values were measured at 490 nm. Control wells containing only medium were used to correct for background absorbance.

For MTT assay, Vero E6 cells seeded at near confluency were incubated with a serial dilution of thapsigargin in a 96-well format. After 24 h, 200 μl MTT mix (DMEM supplemented with 10% FCS containing 250μg/ml tetrazolium bromide, Sigma) was added to each well. Next, cells were incubated for 90-120 min at 37°C and fixed using 3.7% PFA in PBS. The tetrazolium crystals were dissolved by adding 200 μl/well isopropanol and the absorbance at 490 nm was measured using an ELISA reader (BioTek). To determine CC_50_ values, the MTT values were calculated in relation to the untreated control (which was set to 100%). CC_50_ values were then calculated by non-linear regression using GraphPadPrism 5.0 (GraphPad Software).

### ELISA

Sandwich ELISAs from R&D Systems (DuoSet ELISA for human IL-8 (DY208), IL-6 (DY206), CXCL2 DY276-05, CCL20 (DY360)) were used to measure secreted human cytokine /chemokine protein concentrations in cell culture supernatants of HuH7 or MRC-5 cells treated as described in the figure legends. The cell culture supernatants were harvested, centrifuged at 15,000 × g at 4°C for 15 s and stored at −80°C. 100 μl of the supernatants were either used undiluted or were diluted in cell culture medium as follows (HuH7: IL-8 (1:10), CXCL2 (1:3), CCL20 (1:8), MRC-5: IL-8 (1:10), IL-6 (1:20), CXCL2 (1:1.5)) and ELISAs were performed according to the manufacturer’s instructions using serial dilutions of recombinant proteins as standards. All measurements were within the linear range of the standard curve. In some experiments, an IL-1α (10 ng/ml) stimulation for 16 h was used as positive control.

### RNA-seq and bioinformatics

For the data shown in Fig. 1, total RNA was isolated from uninfected and infected cells obtained at 3, 6, 12, 24 h p.i. (or mock infection) using two biological replicates resulting in 32 RNA-seq data sets. RNA was sequenced (with rRNA depletion) using Illumina reagents and an Illumina HiSeq 4000 instrument (single read, 150 bases). Quality control of RNA-seq reads was performed using the FastQC command line tool version 0.11.5 (https://www.bioinformatics.babraham.ac.uk/projects/fastqc/). Reads were aligned using STAR version 2.4.2a (Dobin *et al*, 2013) to an index based on the human genome version hg19. The resulting bam files were imported into R (Team, 2015) (https://www.R-project.org/) and gene-specific read counts based on hg19 UCSC gene annotations were extracted using FeatureCounts from the R subread package version 1.24.2 (Liao *et al*, 2013). Detection of differentially expressed genes was done using DESeq2 version 1.14.1 (Love *et al*, 2014). From the entire data set, only normalized read counts and ratio values for 166 gene IDs assigned to KEGG 04141 were extracted and further analyzed.

### Mass Spectrometry and bioinformatics

Protein extracts were lysed in SDS lysis buffer as described above. Prior to digestion, the SDS containing solution was exchanged to 8 M urea applying the Filter-Aided Sample Preparation (FASP) for proteome analysis protocol using Microcon YM-30 filter devices (Millipore, Cat. MRCF0R030) (Wisniewski, 2018). Cysteines were alkylated with Iodoacetamide and 8 M urea buffer was exchanged to 50 mM ammonium-bicarbonate buffer with a pH of 8.0. Samples were digested within the filter devices by the addition of sequencing grade modified trypsin (Serva) and incubation at 37 °C overnight. Thereafter, the filter-units were transferred to fresh tubes. Peptides were eluted by the addition of 50 μL 0.5 M NaCl solution and centrifugation (14.000 x g for 10 min). After drying the filtrates in a vacuum concentrator (Speed Vac), pellets were resuspended in 25 μL of 0.1% formic acid.

Peptides were desalted and concentrated using Chromabond C18WP spin columns (Macherey-Nagel, Part No. 730522). Finally, peptides were dissolved in 25 μl water with 5% acetonitrile and 0.1% formic acid. The mass spectrometric analysis of the samples was performed using a timsTOF Pro mass spectrometer (Bruker Daltonic). A nanoElute HPLC system (Bruker Daltonics), equipped with an Aurora C18 RP column (25cm x 75μm) filled with 1.7 μm beads (IonOpticks) which was connected online to the mass spectrometer. A portion of approximately 200 ng of peptides corresponding to 2 μl was injected directly on the separation column. Sample loading was performed at a constant pressure of 800 bar. Separation of the tryptic peptides was achieved at 50°C column temperature with the following gradient of water/0.1% formic acid (solvent A) and acetonitrile/0.1% formic acid (solvent B) at a flow rate of 400 nL/min: Linear increase from 2%B to 17%B within 60 minutes, followed by a linear gradient to 25%B within 30 minutes and linear increase to 37% solvent B in additional 10 minutes. Finally B was increased to 95% within 10 minutes and hold for additional 10 minutes. The built-in “DDA PASEF-standard_1.1sec_cycletime” method developed by Bruker Daltonics was used for mass spectrometric measurement. Data analysis was performed using MaxQuant with the Andromeda search engine and Uniprot databases were used for annotating and assigning protein identifiers (Tyanova *et al*, 2016a). Perseus software (versions 1.6.10.50 and 1.6.14.0) was used for further analyses (Tyanova *et al*, 2016b).

For the data shown in Fig. 1, raw data from 47 LC-MS/MS runs (representing two independent experiments and three technical replicates per sample for the 3 h, 6 h, 12 h, 24 h infection time points with the exception of the MERS-CoV 24 h time point which has only five replicates) were mapped to the *Homo sapiens* proteome (uniprot ID UP000005640). The normalized expression values assigned to uninfected HuH7 cells were derived from a total of 59 mock samples representing multiple technical repeats of two biological samples generated at each of the 3 h, 6 h, 12 h, 24 h time points in order to generate a common reference sample for the mean protein expression found in uninfected / untreated HuH7 cells. This mean reference was used to calculate all ratio values. From the entire data set, only protein intensity values for 166 uniprot IDs assigned to KEGG 04141 were extracted, quantile normalized and further analyzed using the software tools described below.

For the data shown in Fig. 4 and 5, raw data from 96 LC-MS/MS runs (representing two independent experiments and three technical replicates per sample) were mapped to *Homo sapiens* (uniprot ID UP000005640 for HuH7 cells), *Chlorocebus sabaeus* (Green monkey, *Cercopithecus sabaeus*, uniprot ID UP000029965 for Vero E6 cells), MERS-CoV (uniprot IDs UP000139997 and UP000171868) or SARS-CoV-2 (uniprot ID UP000464024) peptide sequences. All data sets were processed by MaxQuant version 1.6.10.43 (raw data submission was done with version 1.6.17.0) (Tyanova *et al*., 2016a) including the match between runs option enabled resulting in the identification of 5,372 (from HuH7) or 5,072 (from Vero E6 cells) majority protein IDs. For further quantifications, log_2_-transformed protein intensities were quantile normalized with Perseus (Tyanova *et al*., 2016b) and IDs assigned to contaminants and reverse sequences were omitted. For calculation of ratio values between conditions, the 2 x 6 replicates from each condition were assigned to one analysis group. Differentially expressed proteins were identified from log_2_ transformed normalized protein intensity values by Volcano plot analysis using Perseus functions. All subsequent filtering steps and heatmap representations were performed in Excel 2016 as described in the figure legends. Venn diagrams were created with tools provided at http://bioinformatics.psb.ugent.be/webtools/Venn/. Overrepresentation analyses were done using the gene IDs / gene names of differentially enriched proteins and Metascape software with the express settings (Zhou *et al*., 2019). Protein network data were extracted from the most recent version of STRING (Szklarczyk *et al*., 2019) and visualized with Cytoscape 3.8.0 (Cline *et al*, 2007). Mapping of ratio values on KEEG pathway 04141 was done with Pathview or Pathview Web software (Luo *et al*, 2017).

### Quantification and statistical analysis

Statistical parameters (t-tests, standard variations, confidence intervals, Pearson correlations) were calculated using SigmaPlot 11, GraphPad Prism 5.0 or 8.4.3 or Microsoft Excel 2016 in addition to the software tools mentioned above.

### Data availability

Mass spectrometry raw data are available at https://www.ebi.ac.uk/pride/.

## Notes

### Competing Interest Statement

The authors have declared no competing interest.

## References

Andersen TB, Lopez CQ, Manczak T, Martinez K, Simonsen HT (2015) Thapsigargin--from Thapsia L. to mipsagargin. Molecules 20: 6113–6127

Behnke J, Feige MJ, Hendershot LM (2015) BiP and its nucleotide exchange factors Grp170 and Sil1: mechanisms of action and biological functions. J Mol Biol 427: 1589–1608

Bertolotti A, Zhang Y, Hendershot LM, Harding HP, Ron D (2000) Dynamic interaction of BiP and ER stress transducers in the unfolded-protein response. Nat Cell Biol 2: 326–332

Bojkova D, Klann K, Koch B, Widera M, Krause D, Ciesek S, Cinatl J, Munch C (2020) Proteomics of SARS-CoV-2-infected host cells reveals therapy targets. Nature 583: 469–472

Bouhaddou M, Memon D, Meyer B, White KM, Rezelj VV, Correa Marrero M, Polacco BJ, Melnyk JE, Ulferts S, Kaake RM et al (2020) The Global Phosphorylation Landscape of SARS-CoV-2 Infection. Cell 182: 685–712 e619

Brodsky JL (2012) Cleaning Up: ER-Associated Degradation to the Rescue. Cell 151: 1163–1167

Byun H, Gou YQ, Zook A, Lozano MM, Dudley JP (2014) ERAD and how viruses exploit it. Frontiers in Microbiology 5

Carrara M, Prischi F, Nowak PR, Ali MM (2015) Crystal structures reveal transient PERK luminal domain tetramerization in endoplasmic reticulum stress signaling. EMBO J 34: 1589–1600

Chang TK, Lawrence DA, Lu M, Tan J, Harnoss JM, Marsters SA, Liu P, Sandoval W, Martin SE, Ashkenazi A (2018) Coordination between Two Branches of the Unfolded Protein Response Determines Apoptotic Cell Fate. Mol Cell 71: 629–636 e625

Chen Y, Brandizzi F (2013) IRE1: ER stress sensor and cell fate executor. Trends Cell Biol 23: 547–555

Christianson JC, Olzmann JA, Shaler TA, Sowa ME, Bennett EJ, Richter CM, Tyler RE, Greenblatt EJ, Harper JW, Kopito RR (2012) Defining human ERAD networks through an integrative mapping strategy. Nature Cell Biology 14: 93–U176

Chu H, Dunstl G, Felding J, Baran PS (2018) Divergent synthesis of thapsigargin analogs. Bioorg Med Chem Lett 28: 2705–2707

Chu H, Smith JM, Felding J, Baran PS (2017) Scalable Synthesis of (-)-Thapsigargin. ACS Cent Sci 3: 47–51

Cline MS, Smoot M, Cerami E, Kuchinsky A, Landys N, Workman C, Christmas R, Avila-Campilo I, Creech M, Gross B et al (2007) Integration of biological networks and gene expression data using Cytoscape. NatProtoc 2: 2366–2382

Cui W, Li J, Ron D, Sha B (2011) The structure of the PERK kinase domain suggests the mechanism for its activation. Acta Crystallogr D Biol Crystallogr 67: 423–428

de Wilde AH, Snijder EJ, Kikkert M, van Hemert MJ (2018) Host Factors in Coronavirus Replication. Curr Top Microbiol Immunol 419: 1–42

de Wit E, van Doremalen N, Falzarano D, Munster VJ (2016) SARS and MERS: recent insights into emerging coronaviruses. Nat Rev Microbiol 14: 523–534

Doan NT, Paulsen ES, Sehgal P, Moller JV, Nissen P, Denmeade SR, Isaacs JT, Dionne CA, Christensen SB (2015) Targeting thapsigargin towards tumors. Steroids 97: 2–7

Dobin A, Davis CA, Schlesinger F, Drenkow J, Zaleski C, Jha S, Batut P, Chaisson M, Gingeras TR (2013) STAR: ultrafast universal RNA-seq aligner. Bioinformatics 29: 15–21

Drosten C, Gunther S, Preiser W, van der Werf S, Brodt HR, Becker S, Rabenau H, Panning M, Kolesnikova L, Fouchier RA et al (2003) Identification of a novel coronavirus in patients with severe acute respiratory syndrome. NEngl J Med 348: 1967–1976

Fung TS, Liu DX (2019) Human Coronavirus: Host-Pathogen Interaction. Annu Rev Microbiol 73: 529–557

Gerna G, Campanini G, Rovida F, Percivalle E, Sarasini A, Marchi A, Baldanti F (2006) Genetic variability of human coronavirus OC43-, 229E-, and NL63-like strains and their association with lower respiratory tract infections of hospitalized infants and immunocompromised patients. J Med Virol 78: 938–949

Gorbalenya AE, Baker SC, Baric RS, de Groot RJ, Drosten C, Gulyaeva AA, Haagmans BL, Lauber C, Leontovich AM, Neuman BW et al (2020) The species Severe acute respiratory syndrome-related coronavirus: classifying 2019-nCoV and naming it SARS-CoV-2. Nature Microbiology 5: 536–544

Greenberg SB (2011) Update on rhinovirus and coronavirus infections. Semin Respir Crit Care Med 32: 433–446

Grenga L, Gallais F, Pible O, Gaillard JC, Gouveia D, Batina H, Bazaline N, Ruat S, Culotta K, Miotello G et al (2020) Shotgun proteomics analysis of SARS-CoV-2-infected cells and how it can optimize whole viral particle antigen production for vaccines. Emerg Microbes Infect 9: 1712–1721

Grootjans J, Kaser A, Kaufman RJ, Blumberg RS (2016) The unfolded protein response in immunity and inflammation. Nat Rev Immunol 16: 469–484

Guan Y, Peiris JS, Zheng B, Poon LL, Chan KH, Zeng FY, Chan CW, Chan MN, Chen JD, Chow KY et al (2004) Molecular epidemiology of the novel coronavirus that causes severe acute respiratory syndrome. Lancet 363: 99–104

Halbleib K, Pesek K, Covino R, Hofbauer HF, Wunnicke D, Hanelt I, Hummer G, Ernst R (2017) Activation of the Unfolded Protein Response by Lipid Bilayer Stress. Mol Cell 67: 673–684 e678

Han J, Back SH, Hur J, Lin YH, Gildersleeve R, Shan J, Yuan CL, Krokowski D, Wang S, Hatzoglou M et al (2013) ER-stress-induced transcriptional regulation increases protein synthesis leading to cell death. Nat Cell Biol 15: 481–490

Hetz C, Papa FR (2018) The Unfolded Protein Response and Cell Fate Control. Mol Cell 69: 169–181

Hilton A, Mizzen L, MacIntyre G, Cheley S, Anderson R (1986) Translational control in murine hepatitis virus infection. J Gen Virol 67 (Pt 5): 923–932

Hoffmann E, Thiefes A, Buhrow D, Dittrich-Breiholz O, Schneider H, Resch K, Kracht M (2005) MEK1-dependent delayed expression of Fos-related antigen-1 counteracts c-Fos and p65 NF-kappaB-mediated interleukin-8 transcription in response to cytokines or growth factors. J Biol Chem 280: 9706–9718

Hsu CL, Prasad R, Blackman C, Ng DT (2012) Endoplasmic reticulum stress regulation of the Kar2p/BiP chaperone alleviates proteotoxicity via dual degradation pathways. Mol Biol Cell 23: 630–641

Iwasaki S, Ingolia NT (2017) The Growing Toolbox for Protein Synthesis Studies. Trends Biochem Sci 42: 612–624

Jevsnik M, Ursic T, Zigon N, Lusa L, Krivec U, Petrovec M (2012) Coronavirus infections in hospitalized pediatric patients with acute respiratory tract disease. BMC Infect Dis 12: 365

Jia R, Bonifacino JS (2020) Regulation of LC3B levels by ubiquitination and proteasomal degradation. Autophagy 16: 382–384

Karagoz GE, Acosta-Alvear D, Walter P (2019) The Unfolded Protein Response: Detecting and Responding to Fluctuations in the Protein-Folding Capacity of the Endoplasmic Reticulum. Cold Spring Harb Perspect Biol 11

Kokame K, Agarwala KL, Kato H, Miyata T (2000) Herp, a new ubiquitin-like membrane protein induced by endoplasmic reticulum stress. J Biol Chem 275: 32846–32853

Kopp MC, Larburu N, Durairaj V, Adams CJ, Ali MMU (2019) UPR proteins IRE1 and PERK switch BiP from chaperone to ER stress sensor. Nat Struct Mol Biol 26: 1053–1062

Lee ZW, Low YL, Huang S, Wang T, Deng LW (2014) The cystathionine gamma-lyase/hydrogen sulfide system maintains cellular glutathione status. Biochem J 460: 425–435

Leitman J, Shenkman M, Gofman Y, Shtern NO, Ben-Tal N, Hendershot LM, Lederkremer GZ (2014) Herp coordinates compartmentalization and recruitment of HRD1 and misfolded proteins for ERAD. Molecular Biology of the Cell 25: 1050–1060

Leto DE, Morgens DW, Zhang L, Walczak CP, Elias JE, Bassik MC, Kopito RR (2019) Genome-wide CRISPR Analysis Identifies Substrate-Specific Conjugation Modules in ER-Associated Degradation. Mol Cell 73: 377–389 e311

Li WW, Alexandre S, Cao X, Lee AS (1993) Transactivation of the grp78 promoter by Ca2+ depletion. A comparative analysis with A23187 and the endoplasmic reticulum Ca(2+)-ATPase inhibitor thapsigargin. J Biol Chem 268: 12003–12009

Liao Y, Smyth GK, Shi W (2013) The Subread aligner: fast, accurate and scalable read mapping by seed-and-vote. Nucleic Acids Res 41: e108

Liao Y, Wang X, Huang M, Tam JP, Liu DX (2011) Regulation of the p38 mitogen-activated protein kinase and dual-specificity phosphatase 1 feedback loop modulates the induction of interleukin 6 and 8 in cells infected with coronavirus infectious bronchitis virus. Virology 420: 106–116

Lopez CQ, Corral P, Lorrain-Lorrette B, Martinez-Swatson K, Michoux F, Simonsen HT (2018) Use of a temporary immersion bioreactor system for the sustainable production of thapsigargin in shoot cultures of Thapsia garganica. Plant Methods 14: 79

Love MI, Huber W, Anders S (2014) Moderated estimation of fold change and dispersion for RNA-seq data with DESeq2. Genome Biol 15: 550

Luo W, Pant G, Bhavnasi YK, Blanchard SG, Jr., Brouwer C (2017) Pathview Web: user friendly pathway visualization and data integration. Nucleic Acids Res 45: W501–W508

Ma YJ, Hendershot LM (2004) Herp is dually regulated by both the endoplasmic reticulum stress-specific branch of the unfolded protein response and a branch that is shared with other cellular stress pathways. Journal of Biological Chemistry 279: 13792–13799

Mahalingam D, Peguero J, Cen P, Arora SP, Sarantopoulos J, Rowe J, Allgood V, Tubb B, Campos L (2019) A Phase II, Multicenter, Single-Arm Study of Mipsagargin (G-202) as a Second-Line Therapy Following Sorafenib for Adult Patients with Progressive Advanced Hepatocellular Carcinoma. Cancers (Basel) 11

Mahalingam D, Wilding G, Denmeade S, Sarantopoulas J, Cosgrove D, Cetnar J, Azad N, Bruce J, Kurman M, Allgood VE et al (2016) Mipsagargin, a novel thapsigargin-based PSMA-activated prodrug: results of a first-in-man phase I clinical trial in patients with refractory, advanced or metastatic solid tumours. Br J Cancer 114: 986–994

Maier HJ, Britton P (2012) Involvement of autophagy in coronavirus replication. Viruses 4: 3440–3451

Marder W, McCune WJ (2007) Advances in immunosuppressive therapy. Semin Respir Crit Care Med 28: 398–417

Mehta P, McAuley DF, Brown M, Sanchez E, Tattersall RS, Manson JJ, Hlh Across Speciality Collaboration UK (2020) COVID-19: consider cytokine storm syndromes and immunosuppression. Lancet 395: 1033–1034

Mizutani T, Fukushi S, Saijo M, Kurane I, Morikawa S (2005) JNK and PI3k/Akt signaling pathways are required for establishing persistent SARS-CoV infection in Vero E6 cells. Biochim Biophys Acta 1741: 4–10

Mun K, Punga T (2019) Cellular Zinc Finger Protein 622 Hinders Human Adenovirus Lytic Growth and Limits Binding of the Viral pVII Protein to Virus DNA. J Virol 93

Mungrue IN, Pagnon J, Kohannim O, Gargalovic PS, Lusis AJ (2009) CHAC1/MGC4504 is a novel proapoptotic component of the unfolded protein response, downstream of the ATF4-ATF3-CHOP cascade. J Immunol 182: 466–476

Nakabayashi H, Taketa K, Miyano K, Yamane T, Sato J (1982) Growth of human hepatoma cells lines with differentiated functions in chemically defined medium. Cancer Res 42: 3858–3863

Nicholson KG, Kent J, Hammersley V, Cancio E (1997) Acute viral infections of upper respiratory tract in elderly people living in the community: comparative, prospective, population based study of disease burden. Bmj 315: 1060–1064

Noack J, Bernasconi R, Molinari M (2014) How Viruses Hijack the ERAD Tuning Machinery. Journal of Virology 88: 10272–10275

Okuda-Shimizu Y, Hendershot LM (2007) Characterization of an ERAD pathway for nonglycosylated BiP substrates, which require Herp. Mol Cell 28: 544–554

Oslowski CM, Urano F (2011) Measuring ER stress and the unfolded protein response using mammalian tissue culture system. Methods Enzymol 490: 71–92

Patkar SA, Rasmussen U, Diamant B (1979) On the mechanism of histamine release induced by thapsigargin from Thapsia garganica L. Agents Actions 9: 53–57

Pobre KFR, Poet GJ, Hendershot LM (2019) The endoplasmic reticulum (ER) chaperone BiP is a master regulator of ER functions: Getting by with a little help from ERdj friends. J Biol Chem 294: 2098–2108

Poppe M, Wittig S, Jurida L, Bartkuhn M, Wilhelm J, Muller H, Beuerlein K, Karl N, Bhuju S, Ziebuhr J et al (2017) The NF-kappaB-dependent and -independent transcriptome and chromatin landscapes of human coronavirus 229E-infected cells. PLoS Pathog 13: e1006286

Rota PA, Oberste MS, Monroe SS, Nix WA, Campagnoli R, Icenogle JP, Penaranda S, Bankamp B, Maher K, Chen MH et al (2003) Characterization of a novel coronavirus associated with severe acute respiratory syndrome. Science 300: 1394–1399

Sehgal P, Szalai P, Olesen C, Praetorius HA, Nissen P, Christensen SB, Engedal N, Moller JV (2017) Inhibition of the sarco/endoplasmic reticulum (ER) Ca(2+)-ATPase by thapsigargin analogs induces cell death via ER Ca(2+) depletion and the unfolded protein response. J Biol Chem 292: 19656–19673

Sepulveda D, Rojas-Rivera D, Rodriguez DA, Groenendyk J, Kohler A, Lebeaupin C, Ito S, Urra H, Carreras-Sureda A, Hazari Y et al (2018) Interactome Screening Identifies the ER Luminal Chaperone Hsp47 as a Regulator of the Unfolded Protein Response Transducer IRE1alpha. Mol Cell 69: 238–252 e237

Snijder EJ, Decroly E, Ziebuhr J (2016) The Nonstructural Proteins Directing Coronavirus RNA Synthesis and Processing. Adv Virus Res 96: 59–126

Snijder EJ, Limpens R, de Wilde AH, de Jong AWM, Zevenhoven-Dobbe JC, Maier HJ, Faas F, Koster AJ, Barcena M (2020) A unifying structural and functional model of the coronavirus replication organelle: Tracking down RNA synthesis. PLoS Biol 18: e3000715

Stukalov A, Girault V, Grass V, Bergant V, Karayel O, Urban C, Haas DA, Huang Y, Oubraham L, Wang A et al (2020) Multi-level proteomics reveals host-perturbation strategies of SARS-CoV-2 and SARS-CoV. bioRxiv

Sun S, Shi G, Sha H, Ji Y, Han X, Shu X, Ma H, Inoue T, Gao B, Kim H et al (2015) IRE1alpha is an endogenous substrate of endoplasmic-reticulum-associated degradation. Nat Cell Biol 17: 1546–1555

Szklarczyk D, Gable AL, Lyon D, Junge A, Wyder S, Huerta-Cepas J, Simonovic M, Doncheva NT, Morris JH, Bork P et al (2019) STRING v11: protein-protein association networks with increased coverage, supporting functional discovery in genome-wide experimental datasets. Nucleic Acids Res 47: D607–D613

Team RC, 2015. R: A Language and Environment for Statistical Computing. R Foundation for Statistical Computing, Vienna, Austria.

Tombal B, Weeraratna AT, Denmeade SR, Isaacs JT (2000) Thapsigargin induces a calmodulin/calcineurin-dependent apoptotic cascade responsible for the death of prostatic cancer cells. Prostate 43: 303–317

Tyanova S, Temu T, Cox J (2016a) The MaxQuant computational platform for mass spectrometry-based shotgun proteomics. Nat Protoc 11: 2301–2319

Tyanova S, Temu T, Sinitcyn P, Carlson A, Hein MY, Geiger T, Mann M, Cox J (2016b) The Perseus computational platform for comprehensive analysis of (prote)omics data. Nat Methods 13: 731–740

Urra H, Hetz C (2017) Fine-tuning PERK signaling to control cell fate under stress. Nat Struct Mol Biol 24: 789–790

Wang J, Lee J, Liem D, Ping P (2017) HSPA5 Gene encoding Hsp70 chaperone BiP in the endoplasmic reticulum. Gene 618: 14–23

Wang M, Kaufman RJ (2016) Protein misfolding in the endoplasmic reticulum as a conduit to human disease. Nature 529: 326–335

Wei Y, Meng M, Tian Z, Xie F, Yin Q, Dai C, Wang J, Zhang Q, Liu Y, Liu C et al (2019) Pharmacological preconditioning with the cellular stress inducer thapsigargin protects against experimental sepsis. Pharmacol Res 141: 114–122

Wisniewski JR (2018) Filter-Aided Sample Preparation for Proteome Analysis. Methods Mol Biol 1841: 3–10

Wu H, Ng BS, Thibault G (2014) Endoplasmic reticulum stress response in yeast and humans. Biosci Rep 34

Zaki AM, van Boheemen S, Bestebroer TM, Osterhaus AD, Fouchier RA (2012) Isolation of a novel coronavirus from a man with pneumonia in Saudi Arabia. N Engl J Med 367: 1814–1820

Zhou P, Yang XL, Wang XG, Hu B, Zhang L, Zhang W, Si HR, Zhu Y, Li B, Huang CL et al (2020) A pneumonia outbreak associated with a new coronavirus of probable bat origin. Nature 579: 270–273

Zhou Y, Zhou B, Pache L, Chang M, Khodabakhshi AH, Tanaseichuk O, Benner C, Chanda SK (2019) Metascape provides a biologist-oriented resource for the analysis of systems-level datasets. Nat Commun 10: 1523

Zhu GY, Lee AS (2015) Role of the Unfolded Protein Response, GRP78 and GRP94 in Organ Homeostasis. J Cell Physiol 230: 1413–1420

Zhu N, Zhang D, Wang W, Li X, Yang B, Song J, Zhao X, Huang B, Shi W, Lu R et al (2020) A Novel Coronavirus from Patients with Pneumonia in China, 2019. N Engl J Med

Ziebuhr J, Siddell SG (1999) Processing of the human coronavirus 229E replicase polyproteins by the virus-encoded 3C-like proteinase: identification of proteolytic products and cleavage sites common to pp1a and pp1ab. J Virol 73: 177–185

